# Embodiment in episodic memory through premotor-hippocampal coupling

**DOI:** 10.1101/2023.09.23.559108

**Authors:** Nathalie Heidi Meyer, Baptiste Gauthier, Sara Stampacchia, Juliette Boscheron, Mariana Babo Rebelo, Jevita Potheegadoo, Bruno Herbelin, Florian Lance, Vincent Alvarez, Elizabeth Franc, Fabienne Esposito, Marilia Morais Lacerda, Olaf Blanke

## Abstract

Episodic memory (EM) allows us to remember and relieve past events and experiences, depending on cortical-hippocampal reinstatement involved during encoding. Although it has been claimed that EM is fundamental to establish a sense of self across time, this has never been shown experimentally. Here we combine immersive virtual reality and fMRI and report stronger hippocampal reinstatement for scenes encoded under preserved sense of self, reflecting later recall performance. We further link the sense of self to EM showing that hippocampal reinstatement is coupled with reinstatement in premotor cortex, a key sense of self region. We extend these findings in a severe amnesic patient (caused by bilateral hippocampal damage), whose memory and re-experiencing lacked the normal dependency on the sense of self. Premotor-hippocampal coupling in EM describes how the self at encoding is neurally reinstated during the retrieval of past episodes, enabling a sense of self across time.

**Teaser:** Premotor-hippocampal coupling reveals how the self is reinstated when retrieving past episodes.

## Introduction

Episodic memory (EM) refers to a form of long-term declarative memory associated with the recall of the sensory details of an event (*1*, *2*). EM retrieval is linked to the event’s encoding context and is modulated by various parameters such as the emotional state of the observer and the sensory stimuli of the event during encoding (i.e., visual, auditory, or olfactory cues) (*3–5*, *5*, *6*). A key neural mechanism in EM is the reactivation of brain regions involved at encoding during retrieval, a process which has been called reinstatement and has been observed in several cortical areas (*7–10*). For example, studies reported the activation of primary and extrastriate visual cortex during the retrieval of events containing visual stimuli as well as the reactivation of auditory brain regions during retrieval of events containing auditory stimuli (*6*, *11*). Reinstatement has also been observed in the medial temporal lobe (i.e., in hippocampus, parahippocampus, perirhinal cortex) (*12–15*). Moreover, such hippocampal reinstatement has been associated with memory performance: hippocampal reinstatement was stronger for correctly compared to incorrectly retrieved items (*15*) and was more similar to hippocampal activity at encoding when participants successfully retrieved items. Accordingly, it has been proposed that the hippocampus is a key node necessary for EM recollection, mediating the retrieval of sensory information stored in respective sensory cortical regions (*16–18*), further supported by findings that hippocampal activity at encoding (*19*, *20*) and retrieval (*9*, *15*, *21*, *22*) predicts the reinstatement of cortical areas at retrieval.

However, the sensory context of the *observer’s body* at encoding and its potential reinstatement during the retrieval process has only received scant attention. Thus, it is not known whether sensory bodily inputs - such as tactile, proprioceptive, or vestibular stimuli and their integration with motor signals of the observer’s body at encoding and retrieval - impact EM. This neglect is surprising because the body provides a rich set of sensory-motor inputs during encoding and may provide cues that aid memory formation for visual and auditory stimuli. The few studies that have been carried out revealed that congruent body posture between encoding and retrieval facilitates retrieval of words (*23*) and personal events (*24*). However, none of these studies investigated whether neural reinstatement, as described for the visual and auditory context (*6*, *11*), applies to the bodily sensory context.

Beyond their importance in body representation, certain sensory bodily signals of the observer’s body are also critical for a bodily form of self-consciousness, termed bodily self-consciousness (BSC; 25–28). BSC is based on the integration of multisensory bodily inputs and motor signals (*29–35*) and includes the sense of agency (SoA), body ownership, the first-person perspective (1PP), and self-location (*25*, *27*). Specific components of BSC, such as self-location and 1PP can be altered experimentally using virtual reality (VR), for example by exposing participants to conflicting multisensory visuotactile or visuomotor stimulation (*36*, *37*) from either a first-person (*34*, *38*) or a third-person perspective (3PP) (*34*, *39*, *40*). Visuomotor and perspectival incongruencies have also been shown to modulate SoA (*41–45*). However, despite prominent proposals that self-consciousness is an essential part of EM, as argued by Endel Tulving (*1*, *46*), the impact of experimental alterations of BSC during encoding on later retrieval processes has only recently been investigated. Thus, behavioral evidence demonstrated that the modulation of BSC, using conflicting multisensory and sensorimotor stimulation, influences EM and spatial memory (*47–52*). Although not focusing on BSC framework, the work of St. Jacques and colleagues provided preliminary evidence that BSC may impact EM by showing that the retrieval of an event from a 3PP (compared to 1PP) led to poorer recollection of the sensory and perceptual details experienced at encoding, characterized at the neural level by changes in posterior parietal regions (*47*, *48*, *51–53*). More recently, other studies showed that events seen from a natural 1PP (*54–57*) or with higher body ownership (61), during encoding, were associated with more vivid memories and better memory performance compared to events encoded from the 3PP or without a body view (*54–57*). Collectively, these studies suggest that the modulation of BSC during the encoding of an event affects the later retrieval of that event.

Even fewer studies investigated the underlying brain mechanisms (*54*, *57*, *58*). Such research recorded brain activity either only during encoding or only during retrieval, and focused on the hippocampus and medial temporal lobe structures, thus leaving out the investigation of the coupling of the medial temporal lobe regions with BSC-sensitive areas in fronto-parietal cortex (*54*, *57*, *58*). For example, one study related hippocampal activity during the retrieval of an event to the vividness of the recollection and showed that this depends on the perspective (1PP and 3PP) participants had adopted at encoding (*54*). Another study observed functional connectivity changes between the right hippocampus and the right parahippocampus, depending on whether participants saw their body from their habitual perspective (1PP) or not, again during encoding (*57*). Therefore, it is currently unknown how the neural networks mediating the bodily sensory context and BSC are coupled with EM networks and whether this is based on neural reinstatement.

Here, we investigated how changes in BSC during the encoding of virtual scenes impact EM and neural reinstatement using behavioral experiments, virtual reality (VR), and fMRI. We carried out a series of four VR experiments in a total of 76 healthy subjects as well as in a rare amnesic patient with severe autobiographical memory loss (but preserved BSC). Based on previous work (*55–57*), we designed a VR paradigm and tested EM one hour after the encoding session. For this we combined fully immersive VR with motion tracking and fMRI, allowing us to modulate BSC, by using different levels of visuomotor and perspectival congruency, during the encoding of objects presented in three different virtual scenes. Each scene was associated with a specific experimental condition differing in visuomotor and perspectival congruency (first-person synchronous avatar, first-person asynchronous avatar, and third-person asynchronous avatar). Our participants’ level of BSC modulation was assessed in a separate session (i.e., in a fourth immersive virtual scene). EM was tested one hour after the encoding session using a scene recognition task, critically immersing our participants in the same virtual scenes but without the avatar and the related BSC manipulation. In healthy participants (experiments 1-3), we expected to find higher BSC and better recognition performance for the scenes encoded under visuomotor and perspectival congruency, and higher hippocampal reinstatement in the recognition task for scenes encoded under visuomotor and perspectival congruency, for successful trials. We report that reinstatement-related activity in the hippocampus, activity that we associated with our participants’ performance in scene recognition, is explained by the activation of an independently defined BSC network, including the left dPMC contralateral to the moving right hand. Moreover, such premotor-hippocampal coupling of reinstatement was modulated by visuomotor and perspectival conditions, being stronger for visuomotor and perspectival congruency, linking BSC with the recognition of objects in complex three-dimensional scenes. These data were confirmed and extended by clinical and experimental findings (using the same VR paradigms) in a rare patient, with severe amnesia caused by damage to bilateral hippocampi and adjacent medial temporal structures, who had a normal BSC but was unable to relive and re-experience our complex 3D scenes when she encoded the scenes with visuomotor and perspectival congruency, compatible with a disruption of normal BSC-EM coupling.

## Results

To investigate the effects of BSC on EM, as manipulated by visuomotor and perspectival congruency during encoding, we immersed participants into different 3D virtual environments. The VR paradigm and the visual stimuli were inspired and adapted from previous EM research using immersive VR (*48*, *59*, *60*). Because immersive VR during fMRI is challenging, all participants were first carefully familiarized with the setup and the different virtual environments and were embodied with a virtual body representation. For Experiment 1, we used a head-mounted display and for Experiment 2 MRI-compatible system (see methods). Participants were lying down in a mock MR scanner (Experiment 1) or in the MRI scanner and their movements were tracked with the motion tracking system as described in (*61*). VR paradigms and experimental procedures were implemented using our custom-build system ExVR (ExVR, https://github.com/BlankeLab/ExVR), performing real time control and visual rendering with realistic lighting and shading and, for Experiment 2, synchronizing MRI acquisition with VR experimental data.

We tested memory for complex indoor scenes in three experimental conditions, differing in levels of BSC, manipulated by visuomotor and perspectival congruency. In each condition, participants were immersed and observed the 3D scene and their avatar while moving their right hand. In the first condition, the scene and the avatar were observed from a 1PP and moved synchronously with participants’ upper limb movement (SYNCH1PP; visuomotor and perspectival congruency, no manipulation of BSC; **Fig. 1A**; **Supplementary Video 1**). In the second condition, the avatar was observed from a 1PP but the avatar movement was delayed with respect to the participants’ upper limb movement (ASYNCH1PP; visuomotor mismatch, altered BSC). Finally, in the third condition, participants observed an avatar from a 3PP, and the movement was delayed as well (ASYNCH3PP; visuomotor and perspectival mismatch, strong alteration of BSC). Memory for the scenes was tested one hour after the encoding session in a recognition task where participants were immersed back in the three different virtual scenes (**Fig. 1B**; randomized across participants). Critically, during the recognition task, no avatar or body view was implemented, thus, all scenes, across conditions, were observed from the same location and viewpoint as during encoding, but without any avatar and without manipulation of visuomotor and perspectival congruency. The VR scene was either a scene (**Fig. 1B**) containing the same objects (as shown during the encoding session) or a scene that contained the same number of objects, of which one was changed compared to the encoding session. Healthy participants were asked to report if the scene had changed compared to the encoding session or not. Encoding was incidental in Experiments 1 and 2, that is, participants were not told during encoding that their memory was later tested for the scenes (see methods). In Experiment 1, the task was only behavioral, and in Experiment 2, we also recorded brain activity during encoding, BSC assessment and recognition sessions, with fMRI. Since Experiments 1 and 2 were performed under similar instructions, we combined both samples for behavioral analysis. Because our primary interest was to understand how BSC, as manipulated by visuomotor and perspectival congruency, modulates EM, we compared the synchronous condition seen from the congruent first-person viewpoint (visuomotor and perspectival congruency; SYNCH1PP) with the two conditions in which visuomotor and perspectival congruency was altered (ASYNCH1PP, ASYNCH3PP). BSC and its modulation across the three conditions were assessed using standard questionnaire ratings (see Methods) and, importantly, we used a different complex outdoor scene, to avoid any interference with the encoding of the three scenes used during encoding and recognition sessions (**Fig. 1B**). The sense of agency (SoA) was used to index BSC and its modulation across conditions (see Methods). In the following section, we first describe the behavioral results of Experiment 1 to 3 and then focus on the imaging results of Experiment 2.

**Fig. 1:**
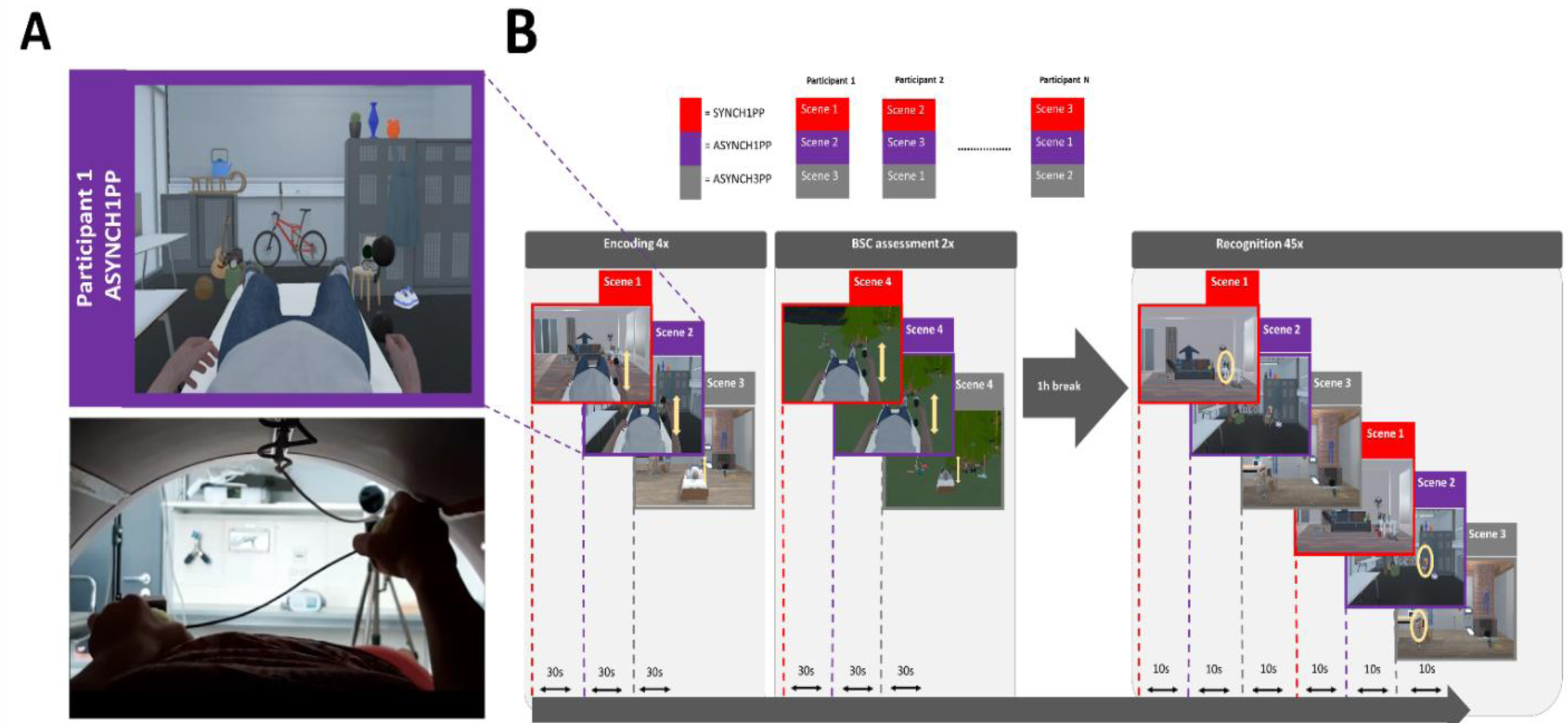
Experimental design: alteration of visuomotor synchrony and perspective in immersive virtual reality. (**A**) Scene snapshot as presented to participants (upper panel) depicts the ASYNCH1PP condition, corresponding to the altered condition of BSC in which the participant was shown the avatar with a first-person perspective (1PP), and with delayed arm movements (lower panel). (**B**) Encoding of three immersive virtual scenes was associated with three different levels of visuomotor and perspectival congruency. One hour after the encoding session, a recognition task assessed scene memory. Note that conditions were attributed solely based on encoding association and the recognition task was always performed without any avatar. BSC sensitivity was tested for each condition in an independent session (with a fourth virtual scene to avoid memory interference). BSC, Bodily self-consciousness.

### Higher SoA during incidental encoding when immersed with visuomotor and perspectival congruency (behavior, Experiment 1 and 2)

To quantify the difference in SoA under the different conditions we applied a linear mixed model for each of the questions asked during the BSC assessment (see Methods). As predicted, SoA was higher in SYNCH1PP as compared to both other conditions with visuomotor and perspectival mismatch (ASYNCH1PP: estimate = -0.07, t = -2.93, p = 0.003; ASYNCH3PP: estimate = -0.07, t = -2.84, p = 0.005; **Fig. 2A**). Ratings were much lower and close to zero for the control questions (**Fig. 2A**; SYNCH1PP compared to ASYNCH1PP: estimate = 0.01, t = 0.94, p = 0.35; SYNCH1PP compared to ASYNCH3PP: estimate = -0.01, t = -1.04, p = 0.3) and differed from the SoA ratings (estimate = -0.47, t = -26.17, p <0.0001). Applying the same approach to the items about body ownership and threat, we found that participants rated their body ownership significantly higher in SYNCH1PP than ASYNCH3PP (estimate = -0.09, t = - 2.9, p = 0.004). Threat was also rated as stronger in SYNCH1PP compared to ASYNCH3PP (estimate = - 0.18, t = -4.45, p <0.0001). The ratings were not significantly different when comparing SYNCH1PP with ASYNCH1PP (body ownership: estimate = 0.01, t = 0.39, p = 0.7; Threat: estimate = - 0.037, t= -0.94, p = 0.35; **Fig. S1A**). There was no difference between Experiment 1 and 2 when adding Experiment as a variable in the model (see Table S1-S4 for the details of the model).

**Fig. 2:**
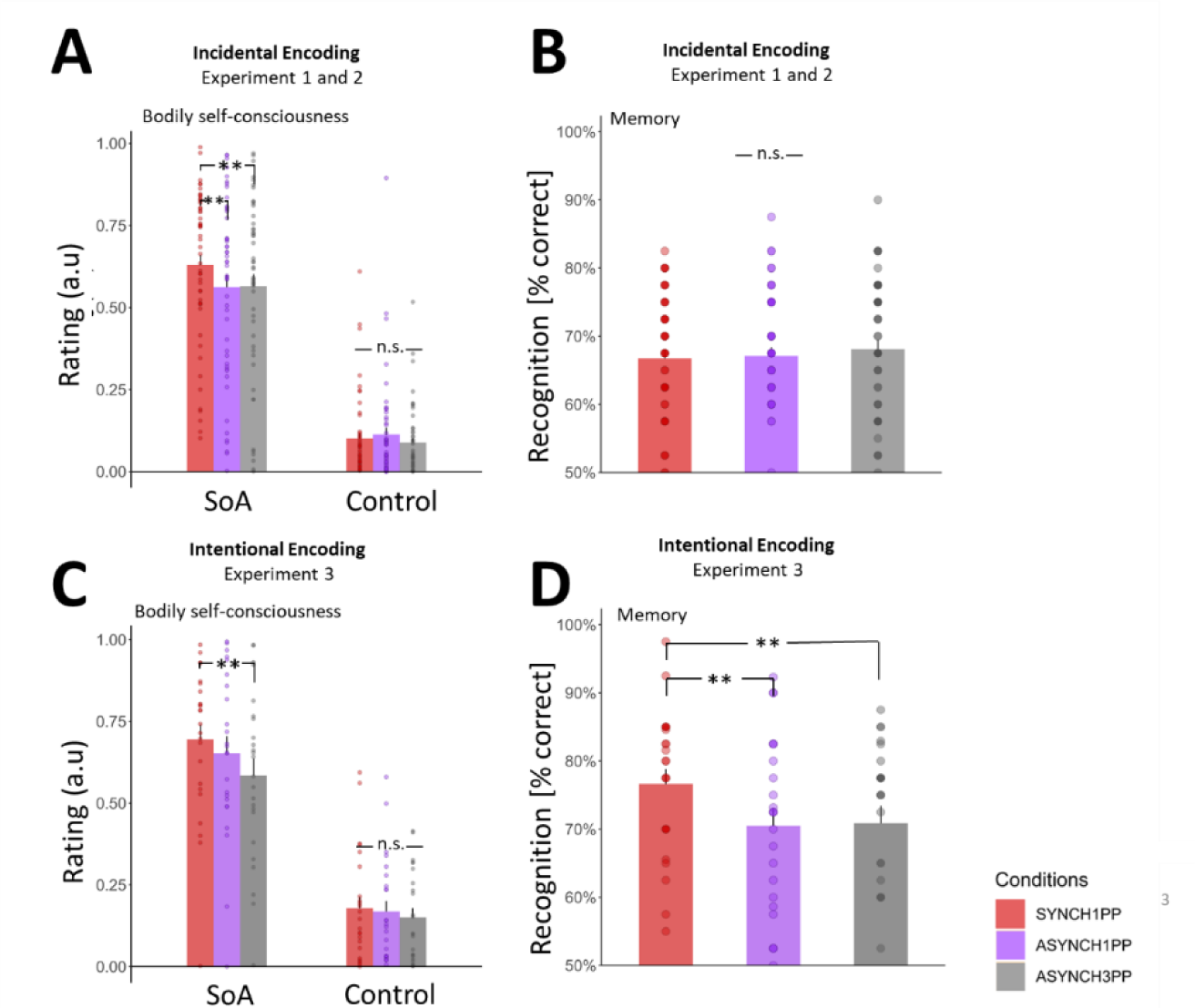
Higher SoA is associated with higher recognition performance under visuomotor and perspectival congruency in intentional encoding. (**A**) When using incidental encoding participants have a higher SoA in SYNCH1PP compared to the two other conditions. Plot of average SoA per (dots) and across (bar) participants. (**B**) During incidental encoding there is no difference in recognition performance between conditions. (**C**) During intentional encoding participants have a higher SoA in SYNCH1PP compared to ASYNCH3PP. (**D)** During intentional encoding, participants better recognized the scenes in SYNCH1PP compared to the other conditions. a.u = arbitrary unit. SoA = Sense of Agency. ** indicates significance level with p-value <0.01 as tested with a linear mixed model using SoA as a dependent variable and conditions as fixed factor.

### Incidental encoding. Visuomotor and perspectival congruency for incidental encoding does not modulate object recognition (behavior, Experiment 1 and 2)

To investigate the effect of visuomotor and perspectival congruency during encoding on recognition performance, we tested participants with a recognition task one hour after the encoding session. Critically, although participants were immersed in the same virtual scenes during the recognition task, they did not see an avatar during the recognition task and there was no manipulation of visuomotor or perspectival congruency. During incidental encoding, participants showed no significant difference in recognition performance (accuracy) between the three experimental conditions (**Fig. 2B**, SYNCH1PP vs. ASYNCH1PP estimate = 0.03, z = 0.4, p = 0.73; SYNCH1PP vs. ASYNCH3PP estimate = 0.06, z = 0.93, p = 0.35). There was no difference in recognition performance between both Experiment 1 and 2 (estimate = -0.02, z = -0.24, p = 0.8, see table S5).

### Intentional encoding. Higher SoA and better recognition performance for intentional encoding when immersed with visuomotor and perspectival congruency (behavior, Experiment 3)

Experiment 3 was similar in all aspects, except that participants were told before the encoding session that their memory for the scenes would be tested subsequently (intentional encoding). As in Experiments 1 and 2, participants’ SoA was higher in SYNCH1PP compared to ASYNCH3PP (**Fig. 2C**; estimate = -0.11, t = -3.16, p = 0.002; the comparison between the SYNCH1PP and the ASYNCH1PP condition was not significant, but similar in direction compared to Experiment 1 and 2 (estimate = -0.04, t = -1.2, p = 0.23). The average ratings for the control items were significantly lower than SoA ratings (estimate = -0.47, t = -17.6, p <0.0001) and not significantly different between conditions (SYNCH1PP compared to ASYNCH1PP: estimate = -0.01, t = -0.56, p = 0.58; SYNCH1PP compared to ASYNCH3PP: estimate = -0.028, t = -1.5, p = 0.13). We also showed higher body ownership and threat ratings in SYNCH1PP versus ASYNCH3PP (body ownership: estimate = -0.2, t= -4.6, p <0.0001; threat: estimate = -0.22, t = -3.13, p = 0.002; **Fig. S1B**), similar to what was found in Experiment 1 and 2. There was no significant difference in body ownership and threat ratings when SYNCH1PP was compared with ASYNCH1PP (body ownership: estimate = -0.06, t = -1.23, p = 0.22; threat: estimate = -0.04, t = -0.5, p = 0.61, Table S7-S10).

For the one-hour delayed recognition task, we found that participants had significantly higher performance in the SYNCH1PP condition compared to ASYNCH1PP (**Fig. 2D**, estimate = - 0.33, z = -3.06, p = 0.002) and ASYNCH3PP (estimate = -0.32, z = -2.95, p = 0.003, Table S11).

To summarize, these three experiments demonstrate that the SYNCH1PP condition induces a higher SoA by exposing participants to objects that were embedded in a 3D VR scene and seen under visuomotor and perspectival congruency during intentional and incidental encoding. Concerning memory, one-hour delayed recognition was boosted for scenes encoded in the SYNCH1PP condition for intentional, but not incidental encoding.

### Better recognition performance for objects presented in the right visual field (behavior, Experiments 1-3)

Because participants moved their right upper limb during encoding, we investigated whether the laterality of objects (right versus left) impacted recognition performance. Such right-handed movements, that were shown in the immersive scenes during the encoding sessions, could have decreased recognition performance due to visual occlusion of the objects placed on the right side or may have, on the contrary, improved recognition performance due to motor facilitation (right-sided upper limb movements) or enhanced attention towards the right side. Results from Experiments 1 and 2 show that participants had better recognition performance for right-sided objects. This was found irrespective of the three conditions (**Fig. S2A**; Table S6; estimate = 0.288, z = 2.028, p = 0.043). We found no such effect of object laterality in Experiment 3 (estimate = 0.037, z= 0.168, p = 0.867 **Fig. S2B**) and no interaction with condition (Table S12), showing that attention or movement-related processes did not differently impact recognition performance in Experiment 3. These data suggest that processes related to attention and/or hand movements may enhance recognition performance for right-sided versus left-sided objects, but only during incidental encoding. Critically, this did not differ between our three experimental conditions in either of the three experiments.

### fMRI (Experiment 2)

Experiment 2 (fMRI, 29 participants) was identical to Experiment 1 (mock scanner, 26 participants), and we recorded fMRI during the three critical periods of the experiment: the encoding session, the BSC assessment session, and the 1-hour delayed recognition session. During all three sessions, participants were exposed to different immersive VR conditions (see **Fig. 1**). We carried out the following fMRI analyses. First, we performed a searchlight representational similarity analysis (RSA) to identify brain areas that were modulated by visuomotor and perspectival congruency and, critically, similarly modulated during both encoding and recognition. Because RSA did not allow us to quantify the *level of similarity* for each condition separately, we computed the encoding-recognition similarity scores (ERS) for each condition and participant and each of the brain regions identified with the RSA. Finally, we investigated if ERS for regions showing significant ERS score differences was associated with participants’ recognition performance.

### RSA analysis. Visuomotor and perspectival congruency impact the reinstatement of encoding activity during recognition

We first explored the condition-dependent reinstatement of encoding activity across the whole brain, to identify reinstatement of brain activity during the encoding and the recognition session, which was similarly modulated by visuomotor and perspectival congruency. For this, we applied a searchlight RSA procedure (**Fig. 3A)** and identified four regions where brain activity showed similar differences that depended on the level of visuomotor and perspectival congruency when comparing encoding with recognition. These regions were left hippocampus, left middle temporal gyrus (MTG), visual cortex, and orbitofrontal cortex (**Fig. 3B**) (Npermutation = 1000, p <0.05, cluster size> 500 voxels).

**Fig. 3:**
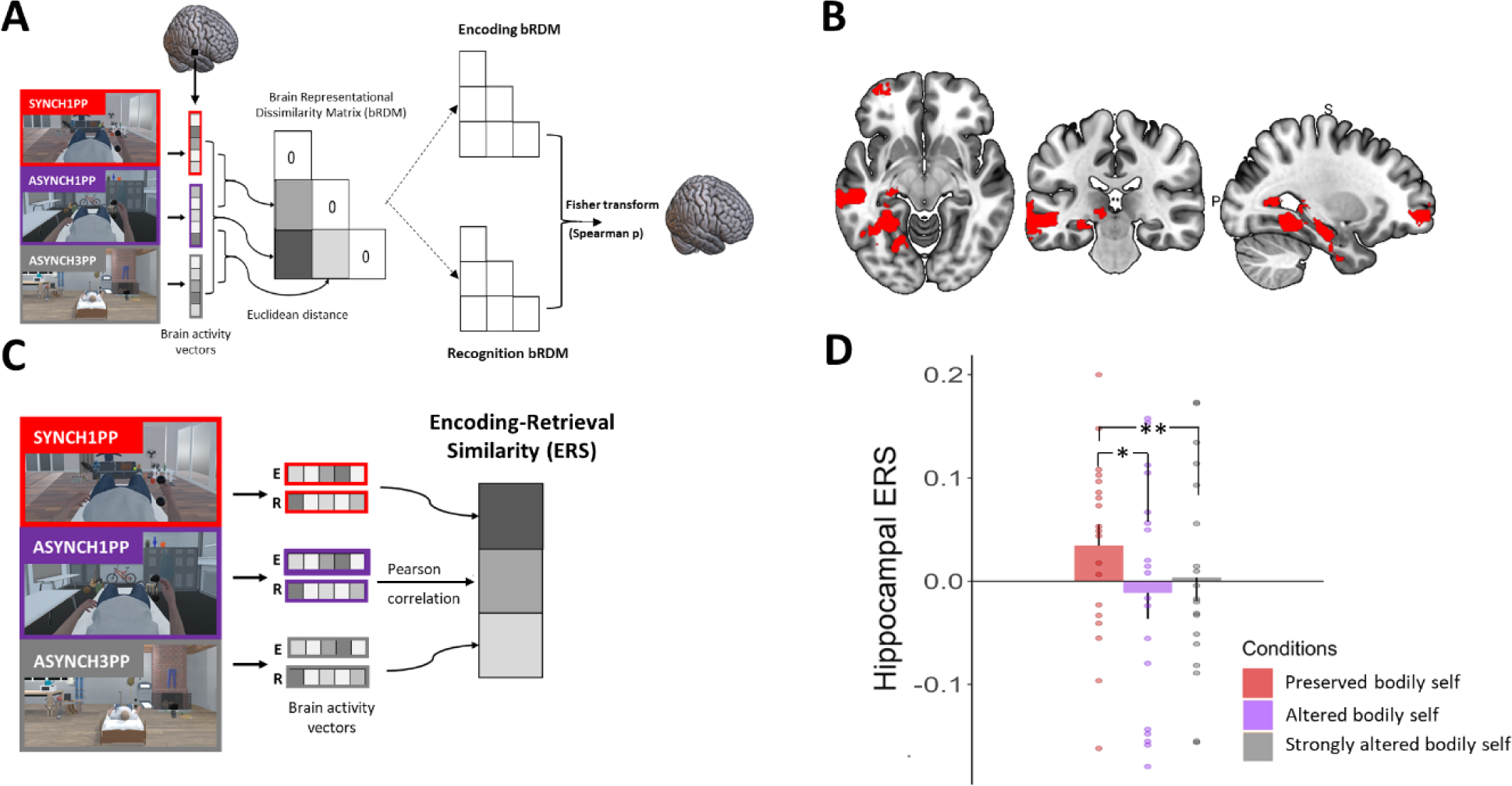
Left hippocampal ERS is higher under visuomotor and perspectival congruency. (**A**) RSA identified brain regions with the same differential pattern between conditions during encoding and recognition sessions in Experiment 2. (**B**) RSA identified four regions: left hippocampus, calcarine cortex, left middle temporal, and left frontal superior orbital gyrus (permutation test, N_permutation_ = 1000, p <0.05, cluster size> 500). (**C**) ERS was computed for the four regions identified by RSA, by applying a Pearson correlation between the voxel activity at encoding and at recognition in Experiment 2. (**D**) Hippocampal ERS is significantly higher under visuomotor and perspectival congruency (SYNCH1PP, red) compared to the two other conditions. * and ** indicates significance level with p-value <0.05 and <0.01 respectively. ERS **=** Encoding recognition similarity score. RSA = Representational Similarity Analysis.

### ERS analysis. Reinstatement in the left hippocampus is higher for visuomotor and perspectival congruency and indexes recognition memory

Although RSA analysis identified four regions that showed a similar pattern of activity between encoding and recognition, it does not provide information about the condition-dependent similarity of activity between encoding and recognition. To further quantify the similarity of activity of these four brain regions between encoding and recognition, we computed their ERS for each of the three conditions separately (*62–64*). This analysis revealed that ERS in the left hippocampus was significantly higher in SYNCH1PP compared to the two conditions with visuomotor and perspectival incongruency (**Fig. 3D**; SYNCH1PP vs. ASYNCH1PP estimate = -0.045, t = -2.57, p = 0.01; SYNCH1PP vs. ASYNCH3PP estimate = -0.045, t = -2.6, p = 0.009, Table S13). There was also a higher ERS in SYNCH1PP compared to ASYNCH1PP in the left MTG (SYNCH1PP vs ASYNCH1PP estimate = -0.048, t = -2.77, p = 0.006, for other comparisons see Table S14). No such ERS differences were observed in visual cortex nor in orbitofrontal cortex (see Table S15-S16 for detailed results).

Based on previous studies showing that hippocampal ERS is a predictor of recognition memory (*65*, *66*), we investigated whether the ERS of the left hippocampus and of the left MTG correlated with participants’ recognition performance. A linear mixed model predicting recognition performance using hippocampal ERS (see Supplementary text and Table S17 for more details about model selection) revealed that ERS of the left hippocampus predicted recognition performance, irrespective of condition (**Fig. 4A**; estimate = 0.29, t = 2.72, p = 0.006, <p_corr_ = 0.0125, Table S18, Supplementary text), such that higher ERS led to better recognition performance. We found a similar relationship between hippocampal ERS and performance in a trial-by-trial model (see Methods), reaching significance only in the condition characterized by visuomotor and perspectival congruency (**Fig. 4B**, Supplementary text and Table S28-33). The same ERS analysis for the left MTG did not detect a significant association with recognition performance (See Table S20).

**Fig. 4:**
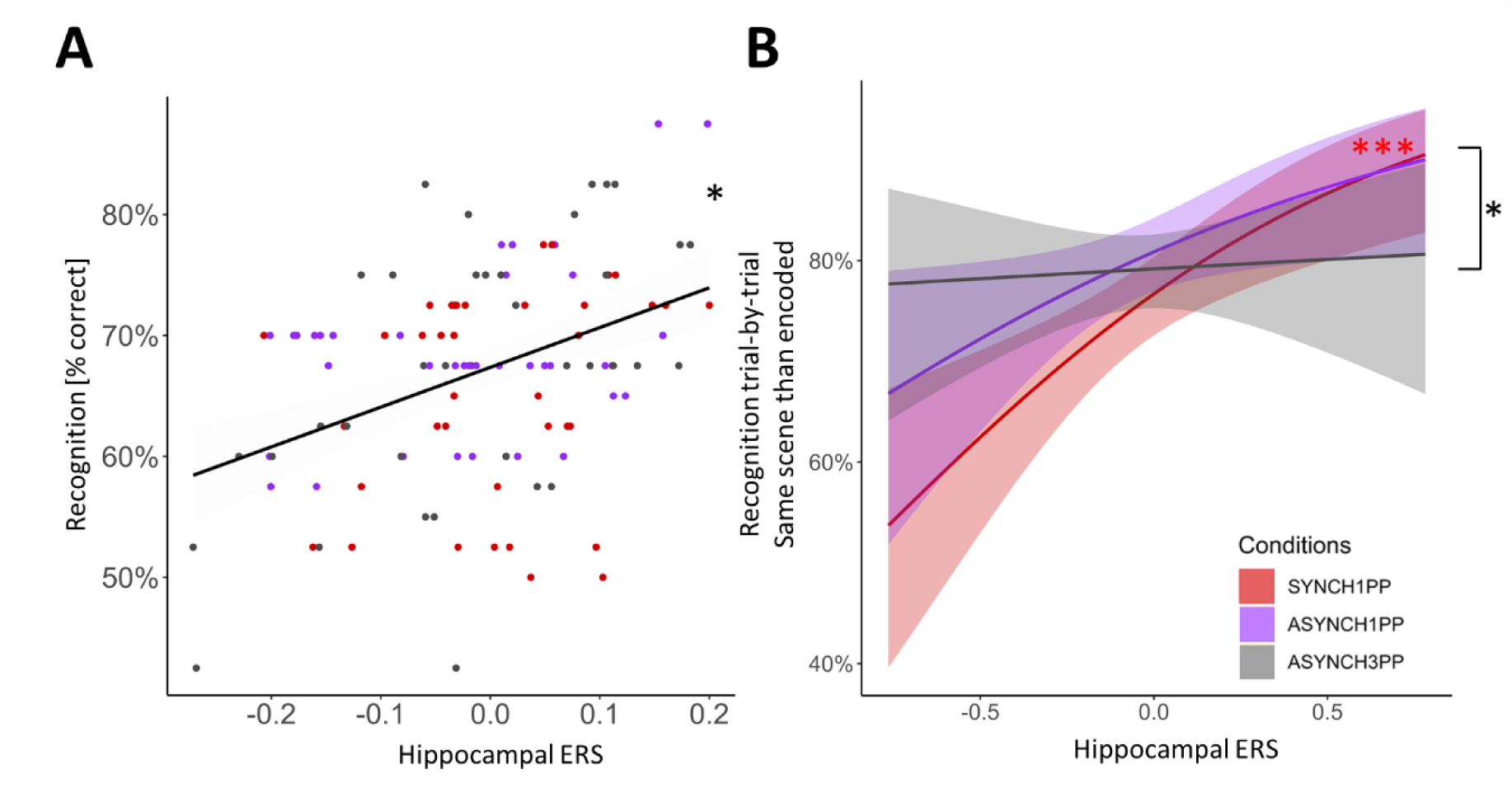
Hippocampal reinstatement of hippocampal encoding activity predicts recognition performance. (**A**) Hippocampal ERS is positively correlated with participants’ recognition performance irrespective of condition (Experiment 2, p = 0.006; p_corrected_ = 0.013; linear mixed model). The mean recognition rate is plotted for each condition (SYNCH1PP: red, ASYNCH1PP: purple, ASYNCH3PP: grey). (**B**) Recognition performance of the original scene is predicted by hippocampal ERS, only in the condition with visuomotor and perspectival congruency. * and *** indicates significance level with p < 0.05 and p <0.0001 (mixed effect logistic regression). ERS **=** Encoding recognition similarity score.

### Hippocampal-neocortical interactions revealed by ERS are modulated by visuomotor and perspectival congruency

Next, we identified brain activity linked to the SoA and investigated whether these SoA regions were associated with the condition-dependent reinstatement of encoding activity. To do so, we first used univariate GLM to identify brain regions modulated by visuomotor and perspectival congruency during the BSC session and then quantified the relationship between ERS of these regions and ERS of the left hippocampus. By contrasting SYNCH1PP with ASYNCH1PP and ASYNCH3PP (SYNCH1PP > ASYNCH1PP+ASYNCH3PP; second level within-subject ANOVA) during the BSC session, we identified a SoA network (**Fig. 5A**) composed of left dorsal premotor cortex (dPMC, MNI coordinate -18, -24, 62) and bilateral supplementary motor area (SMA, MNI coordinate right SMA 4, -4, 55, left SMA -4, -11, 56, Table S19). Post-hoc analysis showed that activity in these regions correlated with participant’s SoA ratings (Supplementary text). Second, we analyzed the interaction between these three brain regions (left dPMC and bilateral SMA) and the left hippocampus (as revealed by RSA and ERS analysis). For this, we applied a linear mixed model investigating how the hippocampal ERS (reflecting recognition performance; dependent variable) was related to the ERS of the left dPMC and bilateral SMA (SoA sensitive regions), for each level of visuomotor and perspectival congruency. This analysis revealed a significant difference in the coupling of the left hippocampal ERS and left dPMC ERS that depended on the experimental condition (**Fig. 5B**; i.e., significant interaction between SYNCH1PP and ASYNCH1PP; ERS dPMC estimate = - 0.19, t = -5.2, p <0.0001, Table S21). These data show that reinstatement in a key memory region (hippocampus) is differently linked with reinstatement in a key SoA region (dPMC), depending on visuomotor and perspectival congruency.

**Fig. 5:**
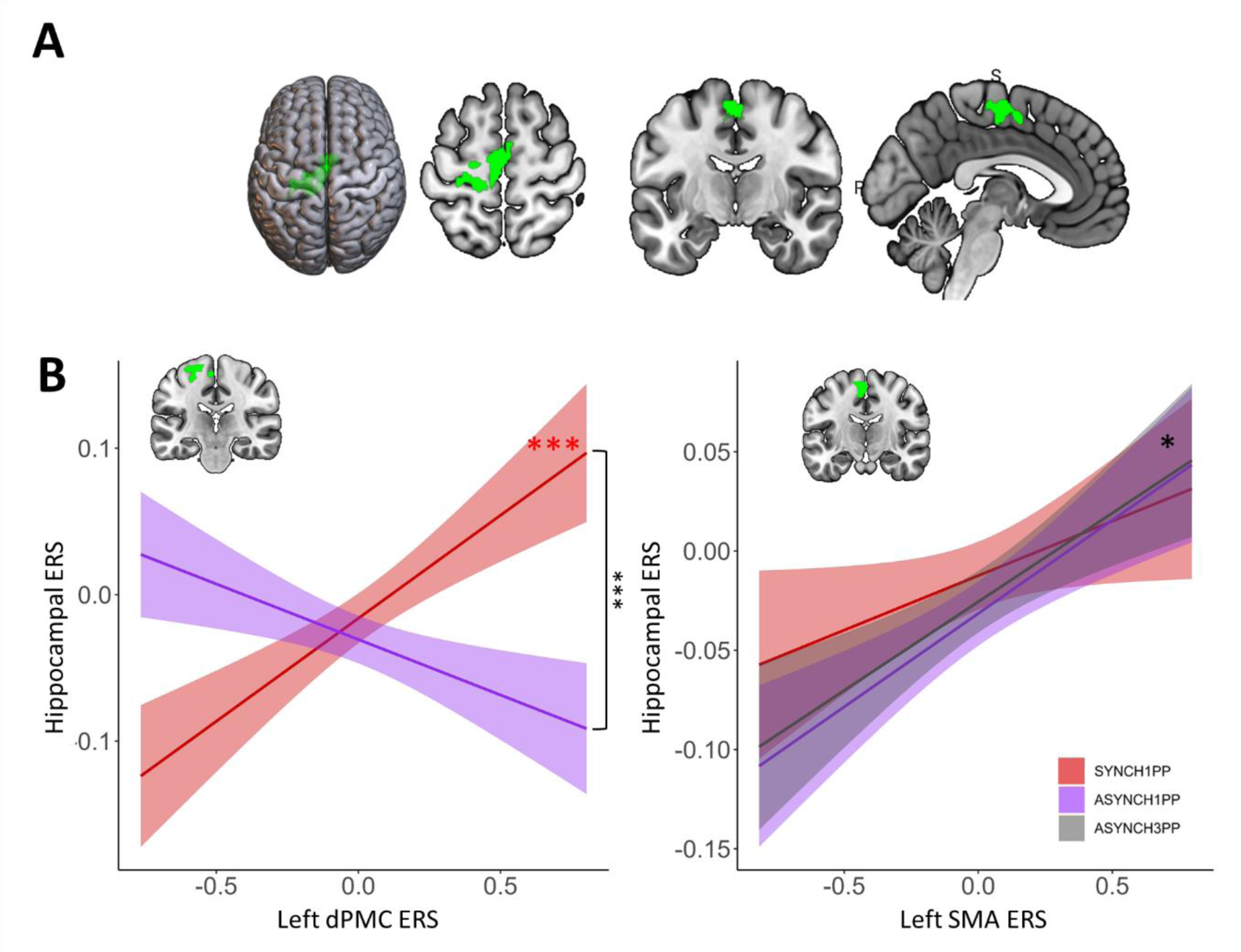
Neural reinstatement of encoding brain activity in SoA-related regions correlates with left hippocampal reinstatement only under visuomotor and perspectival congruency. (**A**) Left dPMC and bilateral SMA activity is higher in the condition with visuomotor and perspectival congruency (SYNCH1PP, red), as compared to the two other conditions. (**B**) Trial-by-trial ERS of the dPMC correlates positively with trial-by-trial hippocampal ERS under visuomotor and perspectival congruency (SYNCH1PP, red) (linear mixed model). Trial-by-trial ERS of the left SMA was found to correlate positively with hippocampal ERS irrespective of condition (p = 0.01, linear mixed model). dPMC = dorsal premotor cortex, SMA = Supplementary motor area, ERS = Encoding recognition similarity score. SoA = Sense of Agency. *, *** indicates significance level with p-value < 0.05 and <0.001 respectively.

This was further extended by post-hoc analysis, revealing that the dPMC ERS was significantly positively related to hippocampal ERS in SYNCH1PP (estimate = 0.18, t = 6.4, p <0.0001), but not significantly related to hippocampal ERS for scenes encoded under visuomotor and perspectival mismatch (for details see Table S22-S23). This shows that higher similarity between encoding and retrieval in the hippocampus (hippocampal ERS) is linked to higher similarity between encoding and retrieval in an SoA sensitive region, dPMC, only in the condition with visuomotor and perspectival congruency and characterized by the highest SoA in the present experiments.

We performed the same analysis for SMA and left hippocampus. Hippocampal ERS was also associated with the left SMA (estimate = 0.06, t = 7.4, p = 0.014, **Fig. 5B**, Table S24). Such hippocampal-SMA coupling was characterized by a positive relation but did not differ between conditions, as found for hippocampal-dPMC coupling. The same analysis applied to the right SMA did not show any coupling with hippocampal ERS (Supplementary text and Table S25).

To summarize, we found that reinstatement-related activity in the left hippocampus, activity that we linked with performance in scene recognition, was systematically related to activity within a cortical SoA network consisting of left dPMC (contralateral to the moving right hand) and bilateral SMA. Whereas hippocampal-SMA coupling was present in all three conditions (reflecting a more general coupling), premotor-hippocampal coupling in the left hemisphere was found for reinstatement-related activity that was stronger under visuomotor and perspectival congruency, suggesting a neural mechanism linking SoA and recognition of objects in complex three-dimensional scenes.

### Amnestic patient with bilateral hippocampal damage is impaired in recognizing objects encoded with visuomotor and perspectival congruency

Would damage to the left hippocampus and its connections with dPMC and SMA, impair the present SoA effects on recognition performance, mediated by visuomotor and perspectival congruency during encoding? We had the unique opportunity to investigate this question in a patient suffering from a severe deficit in autobiographical EM following a fungal brain infection.

The patient is a 62 years old, right-handed woman, who worked as a secretary. She was hospitalized for an epileptic seizure with secondary generalization, followed by several focal epileptic seizures (of temporal origin) with secondary generalizations and status epilepticus (necessitating antiepileptic quadritherapy). The patient showed severe retrograde amnesia (i.e., her daughter’s wedding, holidays, and other important family events) and participated in an intensive neuropsychological rehabilitation program. Despite being able to relearn key facts about her life prior to the infection, the patient is to this day not able to re-experience these key events of her life and her amnesia also extends to new memories following her hospitalization. Repeated neuropsychological examinations revealed a severe EM deficit affecting retrograde events (early childhood to adulthood without temporal gradient) and a moderate anterograde EM deficit. Learning for verbal memory was normal, but deficient for delayed recall (normal after indication). Visuo-spatial memory was at the inferior limit of the norm. There was a mild-to-moderate semantic memory deficit (i.e., public events and celebrities). At the moment of the present investigation (9 months after hospitalization), EM for autobiographical events remained severely deficient with only slight improvements in anterograde EM. A mild executive deficit persisted, but attention was normal.

Seizures were caused by a meningoencephalitis, following a right sphenoidal sinusitis with right sphenoidal bone loss and meningeal contact. Initial MRI (December 2021) showed prominent lesions in the bilateral medial temporal lobe involving both hippocampi, parahippocampi, and amygdalae (**Fig. 6**, **Fig. S3**). Three smaller lesions (all below 1 cm) were also seen in the right middle frontal gyrus, the left inferior parietal gyrus, and right lingual gyrus. Subsequent examinations showed progressive improvement with persistence of lesions in the medial temporal lobe and parietal cortex and disappearance of lesions on MRI in the lingual gyrus and frontal cortex (January 2022). In March 2022 there were no more lesions visible, but the MRI showed bilateral hippocampal atrophy (**Fig. 7A**). This latter finding was further corroborated by volumetric analysis revealing bilateral atrophy affecting both hippocampi (**Fig. 7B, 7D**, Table S26). Hippocampal atrophy affected the four cornu ammoni (CA 1 to 4), the dentate gyrus, the subiculum and the stratum (lacunosum SL; radiatum SR; and moleculare SM). The volumes of amygdala, parahippocampus and entorhinal cortex were in the normal range (Table S26).

**Fig. 6:**
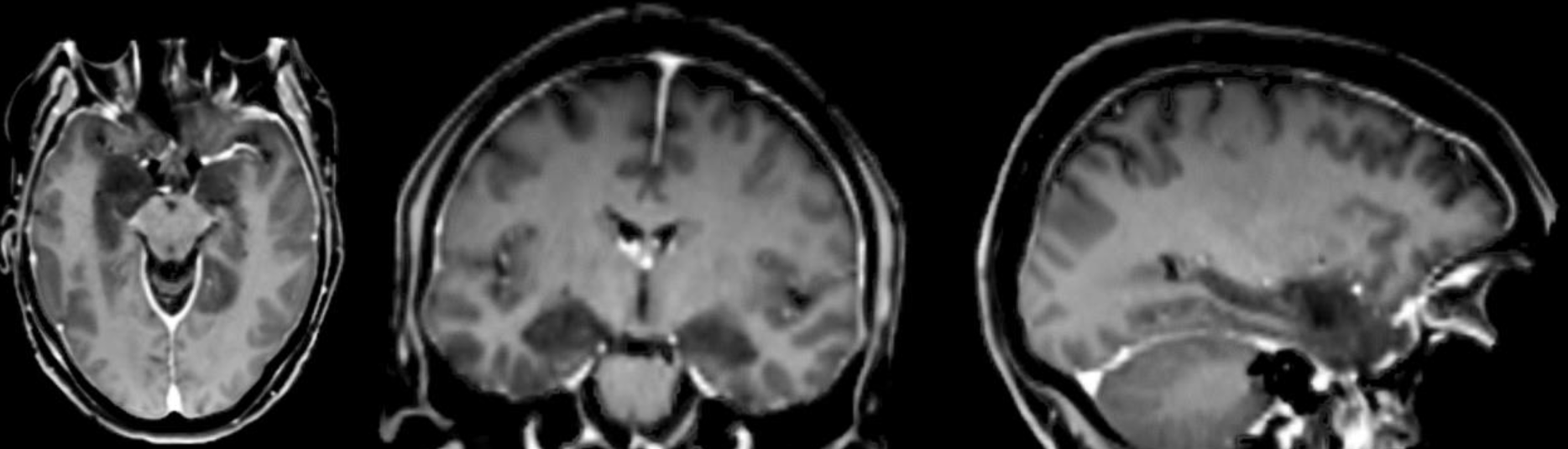
Medial temporal inflamation early during hospitalization of patient. Patient’s structural MRI (T1 MPRAGE, voxel size 1×1×1, 200 slices, 3T MR scanner) during the first week of hospitalization. Dark regions in bilateral medial temporal lobe indicate site of inflammation.

**Fig. 7:**
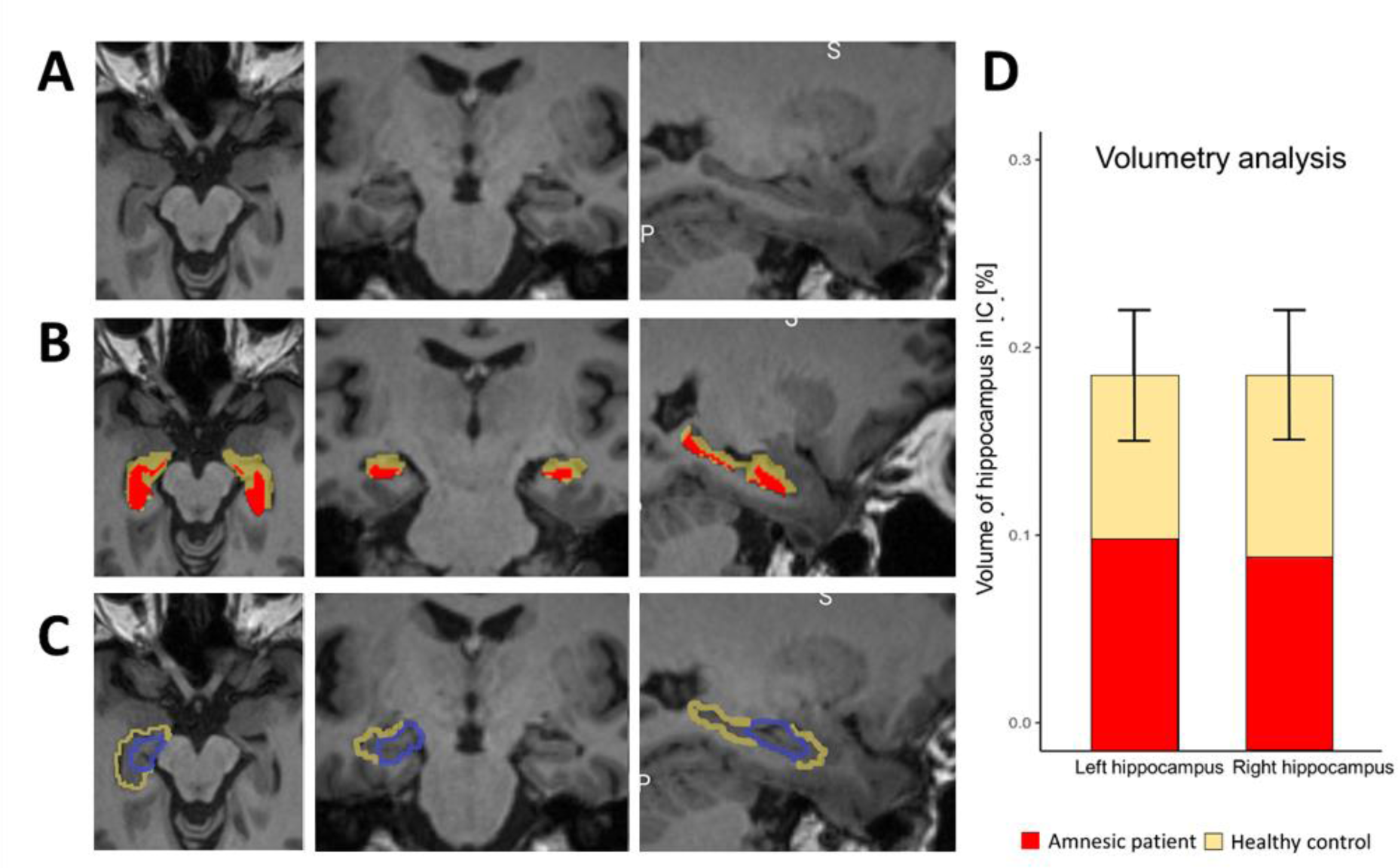
Bilateral hippocampal damage of patient. (**A**) Coronal, axial, and sagital view of the patient’s anatomical MRI eight months after hospitalization (**B**) Patient’s hippocampus (red) is shown with the delineation of a normal hippocampus (yellow), registered in the patient’s native space. (**C**) Hippocampus from the automated anatomical labeling atlas (yellow), and the hippocampal activity, as identified by RSA (blue) in healthy participants, transformed in the patient’s native space overlayed on the patient’s anatomical MRI eight months after hospitalization. (**D**) Volume of the patient’s right and left hippocampi (red bars) compared to the normal range for neurologically healthy age- and gender-matched control (yellow bars). The volume is expressed as the percentage of the hippocampal volume, compared to the total intracranial volume.

The patient’s hippocampal damage involved the left hippocampal region, homologous to the region detected by the present RSA and ERS analysis in Experiment 2, in healthy participants (**Fig. 7C**). Five months after her hospitalization, we tested the patient in the same immersive VR paradigms as tested in healthy participants, adapted to patient comfort (see Supplementary text for details about the adapted setup). We tested the patient with the same immersive VR scenes (as Experiments 1 and 3), in the same three conditions (SYNCH1PP, ASYNCH1PP, ASYNCH3PP), and with the same number of trials. She managed to perform all three sessions: the encoding session, the BSC session, and a one-hour delayed recognition session (see Methods for detail).

As predicted, the patient had preserved SoA, with SoA ratings comparable with those observed in healthy participants in the BSC assessment of Experiments 1-3: she had higher SoA ratings in the SYNCH1PP condition compared to both ASYNCH1PP and ASYNCH3PP conditions (**Fig. 8A)** (Crawford test to compare the patient’s ratings with respect to healthy participants: SYNCH1PP compared to ASYNCH1PP: mean = 0.05, sd ± = 0.16, p < .001; ASYNCH1PP compared to ASYNCH3PP: mean = 0.03, sd ± = 0.16, p = 0.004). The SoA difference between SYNCH1PP and ASYNCH3PP was not significantly different compared to healthy participants but going in the same direction (Crawford test: SYNCH1PP-ASYNCH3PP: mean = 0.08, sd ± = 0.18, p = 0.134). Importantly, the patient’s ratings on control items were low and did not differ from those of healthy participants (**Fig. 8B**) (for a detailed comparison between the patient’s and healthy participants’ SoA ratings see Supplementary text). These data show that the patient was sensitive to our experimental manipulation during encoding, showing a similar modulation of the SoA as healthy participants.

**Fig. 8:**
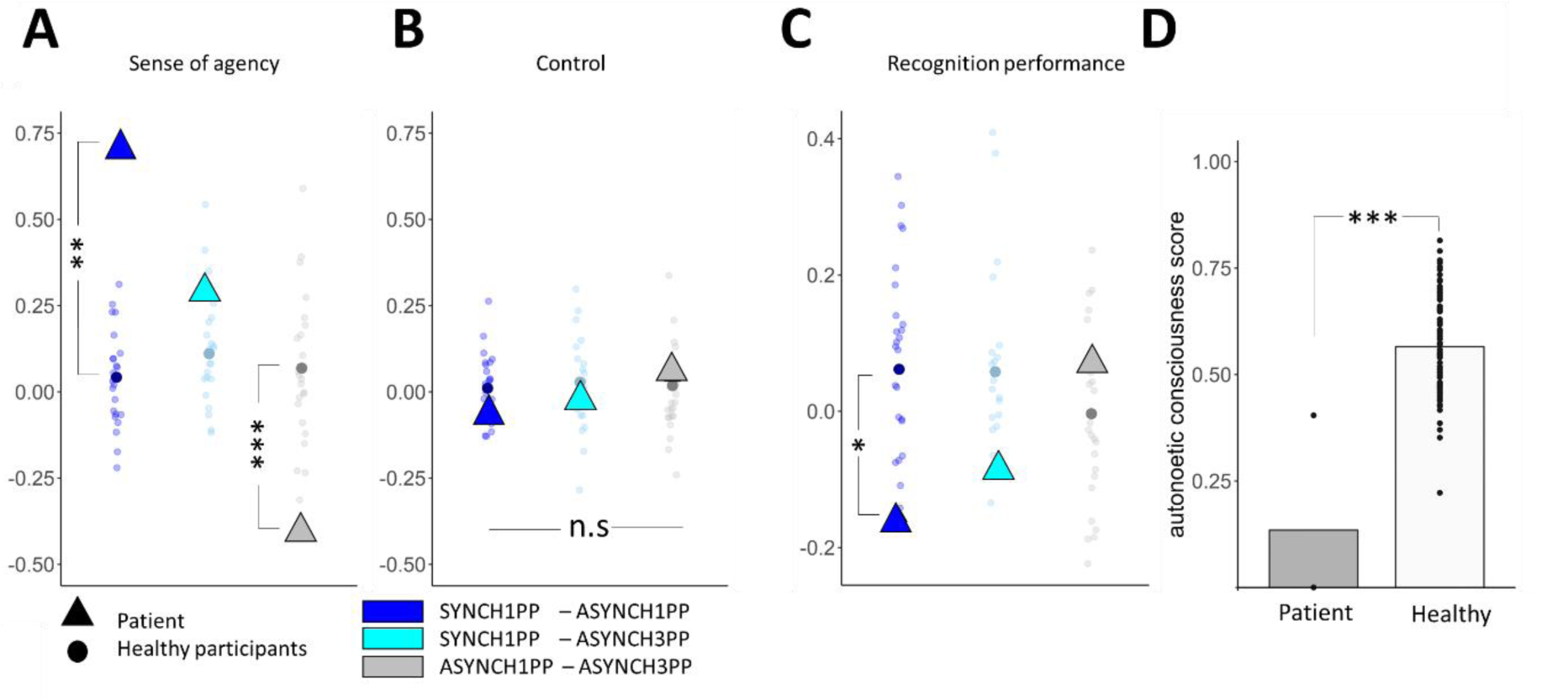
Patient has normal SoA, but abnormally low recognition performance, which does not increase with visuomotor and perspectival congruency. (**A**) Patient has normal SoA (higher SoA for conditions with visuomotor and perspectival congruency versus conditions with visuomotor and perspectival incongruency). The SoA rating difference between SYNCH1PP-ASYNCH1PP (dark blue) and ASYNCH1PP-ASYNCH3PP (grey) are significantly bigger when compared to healthy participants (N = 24) (Crawford test). (**B**) Patient also has normal and low ratings for control questions, which did not differ between conditions and were not different from healthy participants. (**C**) Patient’s recognition performance is lower under visuomotor and perspectival congruency compared to visuomotor and perspectival incongruency differing from healthy participants (colored dots; Experiment 3; Crawford test). (**D**) The patient was able to remember only the scene encoded under the strongest visuomotor and perspectival incongruency (ASYNCH3PP). The autonoetic consciousness of the patient (red) was lower compared to healthy participants (blue; Experiment 3; N= 24) (Crawford test). *, **, and *** indicate significance levels with p-value of <0.05, <0.01 and <0.001, respectively.

The patient was well aware of her memory deficits, for which she had been tested repeatedly during her neuropsychological examinations and memory rehabilitation sessions. Although we initially tested the patient under incidental encoding, she admitted she was expecting this experiment to test her memory. Therefore, we compared her performance with participants who performed the task under intentional encoding instruction (Experiment 3). Inspection of **Fig. 8C** shows that, despite her preserved SoA, she displayed the opposite pattern compared to healthy participants (**Fig. 8C**). She showed the lowest recognition performance in the SYNCH1PP condition (58% correct responses) compared with ASYNCH1PP (73% correct) and ASYNCH3PP (66% correct) conditions. The accuracy difference the patient showed between SYNCH1PP and ASYNCH1PP was significantly different from that observed in healthy participants (Experiment 3) (mean = 0.06, sd ± = 0.11, p = 0.036; the comparison SYNCH1PP-ASYNCH3PP was not significantly different compared to healthy participants (mean = 0.06, sd ± = 0.13, p = 0.148).

To provide further evidence for altered EM, depending on visuomotor and perspectival congruency, we investigated the patient’s autonoetic consciousness, that is, her ability to re-experience the sensory and perceptual details of an event (*1*, *67*, *68*). For this, we asked several questions from the memory characteristics questionnaire (MCQ)(*69*), the episodic autobiographical memory interview (*70*), and the “affected limb intentional feeling questionnaire” (ALEFq, see Supplementary text for more details) (*71*). Although autonoetic consciousness is predominantly tested for autobiographical real-life events, we here tested her autonoetic consciousness for the three virtual 3D scenes into which she was immersed during the encoding session. In particular, we were interested in testing whether her autonoetic consciousness would differ across the three encoding conditions. Based on her lower recognition performance in the SYNCH1PP condition, we predicted that she would indicate lower autonoetic consciousness scores in this condition. Autonoetic consciousness was tested one week after encoding and confirmed this prediction. The patient was able to remember the scene encoded in 3PP (ASYNCH3PP: *“the cabin”*), even without showing her the picture of the scene, and was able to recollect the requested information. However, her overall rating scores in the autonoetic consciousness questionnaire for this scene were still low (**Fig. 8D**). Her memory for the global vividness of the scene was vague (second last choice on a scale from 1 to 7) and global re-experience was low (25%, second last choice on the scale). The averaged ratings significantly differed from the scores of healthy participants as tested with a Crawford test (mean = 0.57, sd = 0.12, p <0.001). More strikingly, she was not able to evoke at all either of the two scenes encoded with a 1PP (i.e., SYNCH1PP: *”the living room”*; ASYNCH1PP: *”the changing room”)*, even after we showed her pictures of these scenes, and was not able to answer and rate the different questions of the ANC questionnaire. The patient reported: *“I can remember seeing the cabin, all in wood, and my arm moving in the scene, but I have never seen this living room nor this changing room”*. The patient showed easier recall for the scene encoded under visuomotor and perspectival mismatch (ASYNCH3PP).

To sum up, damage to bilateral hippocampi and adjacent structures, followed by later atrophy of both hippocampi, led to severe amnesia observed in the present patient. She had a normal SoA, consistent with the fact that dPMC and SMA were not affected by her brain damage. Critically, she remembered fewer objects encoded under visuomotor and perspectival congruency, contrarily to healthy controls. Moreover, she was not able to relive or re-experience the VR scenes that she encoded from a 1PP, but strikingly could only do so for some elements when these were encoded under maximal visuomotor and perspectival mismatch. Further analysis revealed that her brain damage included the left hippocampal region as found in our fMRI analysis in healthy participants (**Fig. 3B** and **Fig. 7C**) and that we linked to visuomotor and perspectival congruency, using RSA and ERS analysis. These clinical-imaging findings further support our previous experimental-imaging findings, that the left hippocampal region and/or its connections with dPMC mediate embodiment effects in EM.

## Discussion

Leveraging the combination of immersive VR and fMRI and accessing brain activity during both encoding and retrieval of episodic memories, we found that (1) the hippocampal reinstatement of encoding-related brain activity during retrieval was higher for scenes encoded with normal visuomotor and perspectival congruency and that (2) such hippocampal reinstatement, critically, reflected recognition performance. We further (3) linked the SoA to hippocampal reinstatement by showing that hippocampal reinstatement was (4) coupled with dPMC reinstatement, a key region of the SoA, that we defined in an independent experimental session and that (5) dPMC activity was also modulated by visuomotor and perspectival congruency, but at encoding. These observations were corroborated and extended in a rare patient with severe amnesia caused by damage and atrophy to bilateral hippocampi, including the hippocampal area critical for reinstatement and dPMC coupling. Although the patient’s SoA was normally modulated by visuomotor and perspectival congruency, she showed worse memory and larger re-experiencing impairments in conditions of normal visuomotor and perspectival congruency. Collectively, these data describe premotor-hippocampal coupling in EM and reveal how the bodily sensory context and the related level of SoA of the observer at encoding is neurally reinstated during the retrieval of past episodes.

### Hippocampal reinstatement reflects recognition performance and the visuomotor and perspectival congruency of the encoded scenes

We found that the hippocampal reinstatement of encoding activity across the experimental conditions was correlated with participants’ recognition performance, consistent with previous work using visual or auditory stimuli (*12*, *15*). Thus, hippocampal reinstatement - reflected by the average hippocampal activity across all trials - correlated with average recognition performance (Fig. 4A). This is consistent with the idea that successful EM retrieval depends on the degree of remobilization of activity observed during encoding (*15*, *19*), i.e. reactivation of the hippocampal engram (*72*). Neural reactivation of the hippocampus during EM retrieval and its link with memory performance has been demonstrated previously (*10*, *19*, *22*, *73*). Yet this earlier work presented single or paired stimuli at encoding (i.e., pictures, word cue (72,61), whereas the present results report hippocampal reinstatement in a richer sensory context with action-embedded 3D scenes using immersive VR in fMRI with incidental encoding, closer to encoding conditions in our everyday life.

These findings were extended by additional trial-by-trial analyses showing that hippocampal reinstatement reflects the successful recognition of the original scene presented during encoding for individual trials and only when encoding was done with visuomotor and perspectival congruency (SYNCH1PP condition, Fig. 4B). These data suggest that the successful recognition of the scene observed at encoding is critically linked to the reactivation of the hippocampal encoding activity during retrieval, extending previous evidence about the hippocampus’ role in pattern separation to discriminate between previously encoded events and new events (*14*, *74–76*).

This further indicates that EM retrieval not only depends on stronger similarity between the encoding and retrieval patterns in the hippocampus, but also on preserved visuomotor and perspectival congruency at encoding. This demonstrates that hippocampal reinstatement depends on the bodily sensory context of the observer during encoding, because only the neural pattern of episodes encoded from a first-person perspective and with synchronous movements helped distinguish between the events presented at encoding and new events at retrieval.

We further show that hippocampal reinstatement of the averaged activity (across all trials) also depends on the level of the SoA at encoding, differing between the three experimental conditions. Critically, we show that the highest level of hippocampal reinstatement was found for scenes encoded under visuomotor and perspectival congruency. This is in line with a recent study linking hippocampal reinstatement with the modulation of body ownership during encoding (*58*), although the latter work did not observe reinstatement using full brain analysis (but rather used a region of interest approach) and did not associate hippocampal reinstatement with EM performance memory or the modulated BSC level.

In addition to EM-related reinstatement in the hippocampus, we also observed reinstatement activity in the left MTG that was further modulated by the SoA level. Although this region has been reported in previous EM studies (using video stimuli) (*8*), MTG reinstatement in the present study was not correlated with recognition performance, showing that reinstatement functionally differs between MTG and hippocampus and that only hippocampal reinstatement reflected SoA as well as EM performance. We did not find any relationship between recognition performance and cortical reinstatement in other regions such as visual cortex, which was detected by RSA analysis. Although several studies provided evidence of remobilization of visual cortex during retrieval (*77*, *78*), our results show that this reinstatement does not reflect visuomotor and perspectival congruency.

We argue that the present data provides evidence for embodied hippocampal reinstatement by showing that the sensorimotor context of the observer’s body at encoding impacts encoding- and retrieval-related hippocampal activity. The finding of embodied hippocampal reinstatement (i.e., reinstatement that depends on visuomotor and perspectival congruency) extends previous reinstatement observations that investigated the visual or auditory information of the encoded scene, to the bodily sensorimotor context of the observer during the encoding of the scene.

### Hippocampal – premotor coupling is stronger in the left dorsal premotor cortex under visuomotor and perspectival congruency

By showing that the dPMC exhibited higher activation under visuomotor and perspectival congruency the present data support the PMC’s well-known role in motor control and motor-related cognition, consistent with previous brain imaging studies on the SoA (38,78–80). We note that the hippocampal reinstatement (that was associated with EM performance and depended on visuomotor-perspectival congruency) as well as the dPMC activity (depending on visuomotor-perspectival congruency) were both found only in the left hemisphere, that is in the hemisphere contralateral to the participants’ right upper limb movements during encoding (*79*), suggesting a further functional link of the bodily sensory context during encoding on premotor and hippocampal activity in the present experiment. Several reinstatement studies have observed left hippocampal reinstatement (*8*, *22*, *54*, *58*). While some studies found that especially left hippocampal activity was linked with memory performance (*8*, *22*) and associated with the perspective during encoding (53), others did not report hemispheric specificity of hippocampal reinstatement (*13*, *62*). Further work is needed to investigate the lateralization of hippocampal activity during reinstatement processes and how this depends on the lateralization of sensory stimuli and movements of the participant.

Several studies have demonstrated that hippocampal activity during encoding is coupled with the reactivation of other cortical regions, during the retrieval process (14,82). In particular, activation of visual cortex has been linked with hippocampal activity, at encoding and retrieval, and it has been suggested that hippocampal reinstatement mediates the reinstatement of cortical areas during retrieval (*14*, *21*). In our study, we found that the reinstatement of the left hippocampus was linked to left dPMC reinstatement, at the single trial level (Fig. 5B). Together with past experimental work, this observation extends the widely accepted theory that the hippocampus indexes sensory information of EM stored in visual and auditory regions (17,18,81) to indexing sensorimotor information stored in dPMC. Thus, the present results show that hippocampal-cortical coupling varies in function of the sensory encoding context and that, in the case of different bodily sensory contexts and SoA levels during encoding, activity in dPMC is coupled with the hippocampal activity. Moreover, strongest premotor-hippocampal coupling was found in the present study for scenes encoded under preserved SoA. Accordingly, our results provide further evidence for embodied reinstatement, showing that not only hippocampal reinstatement, but also the coupling between hippocampus and sensorimotor cortices depends on preserved normal SoA levels, as induced here by visuomotor and perspectival congruency.

We also report bilateral SMA-hippocampal coupling. The SMA is a brain region involved in movement selection and motor preparation and has also been involved in the SoA (38,78–80). Bilateral SMA was more activated under visuomotor and perspectival congruency (independent of the reinstatement analysis) and, critically, characterized by a trial-by trial coupling between left hippocampal reinstatement and SMA reinstatement. However, while the premotor-hippocampal coupling (i.e., with left dPMC) was specific to the condition with preserved visuomotor and perspectival congruency condition, the coupling between the reinstatement of the left SMA and the left hippocampus was independent of our experimental conditions. These data show that SMA-hippocampal coupling also contributes to reinstatement, but that the reinstatement mechanism is a more general one that does not reflect visuomotor and perspectival congruency and the related SoA (compared to the embodied reinstatement that is mediated by premotor-hippocampal coupling).

### Decreased recognition performance under visuomotor and perspectival congruency in an amnesic patient with bilateral hippocampal damage

Severe EM deficits including autobiographical EM have been described previously in several patients with bilateral hippocampal damage (*68*, *80–82*). Here, we investigated the impact of the bodily sensory context and of BSC on EM, using the paradigm of experiments 1-3 in an amnestic patient, with a severe retrograde and a moderate anterograde EM deficit. Her recognition performance in all three conditions was significantly lower as compared to healthy participants in experiments 1 and 2 (Fig. 8C) and so was her autonoetic consciousness (Fig. 8D). Concerning the SoA, the patient had normal levels of BSC and showed normal modulation of her SoA ratings that depended on visuomotor and perspectival congruency, comparable to healthy participants (Fig. 8A) and comparable to previous studies in healthy subjects (*83–86*). Critically, the patient’s EM performance was lowest in conditions when her SoA was increased (i.e., the condition with visuomotor and perspectival congruency) (Fig. 8C), thus showing the opposite behavioral pattern as observed in healthy participants, especially experiment 3. This selective condition-dependent modulation was further confirmed when testing her autonoetic consciousness one week after the encoding session, when the patient was only able to re-experience the scene that she had encoded with the weakest SoA and thus the strongest visuomotor and perspectival incongruency (i.e., ASYNCH3PP) (Fig. 8D).

The present patient suffered from bilateral damage and later atrophy of both hippocampi (Fig. 7A), causing her memory and autonoetic consciousness deficits. Other regions in the medial temporal lobe (i.e, amygdala, entorhinal cortex) were not significantly reduced in volume compared to a healthy age-matched control group. Her lesion overlapped with the left hippocampal region identified in this study (Fig. 7C), but did not involve the left dPMC premotor cortex nor the SMA. This damage changed the way the bodily sensory context and BSC during encoding impact her EM, while keeping her SoA preserved. We argue that the altered modulation of EM by visuo-motor and perspectival congruency is caused either by damage to the left hippocampus or to structural or functional changes in premotor-hippocampal coupling in the present patient. This is compatible with animal work showing that the silencing hippocampal activity at retrieval prevents the reactivation of cortical areas involved during encoding processes (*7*, *87–89*). Although there are many reports of single case patients with amnesia due to lesions in the medial temporal and frontal lobe (*68*, *80*, *82*, *90*), to our knowledge, this study is the first to assess the effect of BSC manipulation in an amnesic patient and therefore provide novel clinical insight into the neural association between BSC and EM.

In conclusion, we report that hippocampal reinstatement of encoding-related brain activity during retrieval was higher for scenes encoded with normal visuomotor and perspectival congruency and reflected recognition performance. We further linked hippocampal reinstatement with dPMC reinstatement, a key region of the SoA. Premotor-hippocampal coupling thus appears as a mechanism for the reinstatement of the bodily sensory context and the SoA during the retrieval of a past episode. These observations were corroborated and extended in a rare patient with severe amnesia caused by damage and atrophy to bilateral hippocampi, including the area of hippocampal reinstatement and coupling. Together, our studyprovides behavioral, imaging and clinical evidence of the involvement of bodily sensory context and BSC in the neural bases of EM.

### Study limitations

This study did not aim to separate the specific mechanisms associated with the first-person perspective or with visuo-motor synchrony, but to provide first evidence into the neural mechanisms linking BSC and EM. Therefore, we tested the effects of graded conditions, from preserved BSC (SYNCH1PP), to moderate (ASYNCH1PP), to strong BSC alterations (ASYNCH3PP). Future studies may investigate the specific effects of perspective and congruency on the behavioral, neural, and clinical mechanisms leading to the present coupling of BSC and EM. We also note that observations in single patients should be regarded with caution. The present neuropsychological-behavioral effects need to be confirmed in future clinical studies in amnesic patients and compared with age-matched healthy control groups.

## Methods

### Experimental Design

All experiments consisted of three separate main sessions. The first session was a memory encoding session, which was followed by an assessment of bodily self-consciousness (BSC; see below). The last session was a recognition session carried out one hour later (**Fig. 1**). We also assessed autonoetic consciousness one week after the encoding session. Before the experiment, all participants underwent a familiarization session with VR for each of the four scenes (the three encoding scenes and the BSC assessment scene) that were used during the task (See Supplementary method). For Experiment 1 to 3, participants were lying in supine position (in a mock replicate of a MR scanner for Experiment 1 and in the MRI for Experiment 2) and equipped with VR headsets. They hold tennis ball in each hand equipped with button responses and reflective marker to track their movement. The following section explain the detail setup for each session and experiment.

### Encoding session

During the encoding session, participants were instructed to keep moving their right upper limb while observing a virtual avatar animated in real-time. Upper limb movements were instructed to occur between two virtual black spheres that were displayed in the visual scene. The black spheres were aligned vertically, to the right of the virtual body, and placed at the level of the avatar’s hip (**Fig. 1**). We manipulated the sense of agency (SoA; 92–94) of participants by exposing them to three different scenes corresponding to the three different experimental conditions, which were characterized by different levels of sensorimotor synchrony. For this, participants were exposed to different levels of visuomotor congruency between the movement of their right upper limb and the shown movements of the avatar’s upper limb in the virtual scene (i.e. 43,95). In the SYNCH1PP condition there was no visuomotor manipulation thus, the virtual avatar was seen from a first-person perspective (1PP) and the right virtual upper limb was moving synchronously to the participant’s upper limb movements (SYNCH1PP). In the ASYNCH1PP condition, the virtual avatar was also seen from a 1PP, but was moving with a visuomotor delay that varied between 800-1000 ms to the movement of the participant (ASYNCH1PP). In a third, control, condition (ASYNCH3PP), the avatar was seen from a third-person perspective (3PP) and was moving with a visuomotor delay that varied between 800-1000 ms with respect to the movement of the participant. For the 3PP the body of the avatar was moved forward in the virtual scene to maintain the same visual angle of all objects in the scene, as compared to the other experimental conditions. Based on previous work on SoA (*96–98*), we expected stronger SoA in the condition with no visuomotor manipulation and naturalistic perspective (SYNCH1PP) compared to the conditions with visuomotor manipulation (ASYNCH1PP and ASYNCH3PP).

Each of the three encoded scenes contained eighteen objects and was associated with a specific experimental condition (SYNCH1PP, ASYNCH1PP, ASYNCH3PP) for each participant. This association between the encoded scene and experimental condition was pseudo-randomized across participants. For each experimental condition, each scene was presented for 30 seconds and repeated four times. An inter-trial interval consisting of a fixation cross appeared for five seconds in between each scene presentation to avoid potential carry-over effects from one condition to another.

Encoding was incidental in Experiments 1, 2 and 4. Thus, participants were not told that they participated in a memory experiment and that their object recognition was going to be tested. Encoding was intentional in Experiment 3 and we instructed the participants to pay close attention to the scene during encoding and told them they would be tested on the scene one hour later. The rest of the experimental design was the same between Experiments 1, 2 and 3.

### BSC assessment

Immediately after the encoding session, participants were immersed in a different outdoor scene containing eighteen new objects to avoid any memory interference with the encoding of the 3 scenes associated with the 3 experimental conditions. They were instructed to perform the same right upper limb movements and were observing the same avatar in SYNCH1PP, ASYNCH1PP, and ASYNCH3PP, but now performed in the outdoor environment (duration 30 seconds). Based on previous BSC work, in Experiments 1, 2 and 3, we also included a response to a threat stimulus directed towards the avatar (i.e. after 30 seconds, an unexpected event consisting in a virtual knife seen as approaching the avatar’s trunk (*49*, *99*).

Participants had to rate their agreement with five statements regarding different aspects of BSC: (Q1) “I felt that I was controlling the virtual body” to rate their SoA toward the movement of the virtual avatar; (Q2) “I felt that the virtual body was mine” to rate their level of body ownership toward the virtual avatar intentionally; (Q3, only in Experiment 1,2 and 3) “I was afraid to be hurt by the knife” to rate their threat response as a proxy of the subjective measure of their BSC as in(*49*). We also included two control statements for experimental bias: (Q4) “I felt like I had more than three bodies” and (Q5) “I felt like the trees were my body”. The five statements were presented successively and in a randomized order. For each statement, a cursor was programmed to move between the two extreme points of the presented agreement scale (between 0 to 1, with an increment of 0.001) at a constant speed. The participant had to stop it at the desired position by a left button press and then validate their response with a right button press. Before validation, the participant was free to retry indefinitely to specify his agreement level by a left click until being satisfied by the answer. The BSC assessment was repeated twice per condition.

### Recognition session

One hour after the encoding session, participants were presented with the encoded scenes again. Participants were exposed to each tested scene for 10 seconds and were then asked to respond yes or no to the question: “Is there any change in the room compared to the first time you saw it?”. They were instructed just before the start of the recognition session that they will have to answer concerning the original scenes seen during the encoding session. Some of these scenes were identical to the encoded scenes (original scene), and other were modified (changed scene). Participants performed 45 trials per condition. Among those 45 trials, 20 trials corresponded to the presentation of the original scene and 20 trials corresponded to a modified version of the original scene in which one single object was changed in either color or shape. There were 5 additional attentional trials in which two to three objects were changing shape, color, or position in the scene. The attentional trials were used to incite the participants to carefully observe the entire scene instead of simply trying to spot a single object change. Attentional trials were not included in the analysis. Importantly, during the recognition session, participants did not move their arms and no avatar was shown to not modulate BSC during the recognition session. In Experiment 1, following the first statement participants had to answer an additional question: “How confident are you about your answer?”. They had to answer using the same cursor-stopping scheme as in the BSC assessment following the encoding session. In Experiments 2,3 and 4, the second question was replaced by a 3-second fixation cross, to reduce experimental time as well as to avoid carry-over effects.

### Autonoetic consciousness session

We also tested the patient and the healthy participants’ autonoetic consciousness for each condition, at one week after the encoding. Autonoetic consciousness was assessed using an association of questions from a well-established questionnaire for a total of 31 questions, including questions from the “Memory characteristic questionnaire” (19 questions *69*), part B of the “Episodic autobiographic memory interview” (EAMI; 8 questions (*70*), two questions from the “affected limb intentional feeling questionnaire” ALFq (*71*) and one additional question related to our research question (“I remember the movement and gesture I was doing with my body during the event”, ordinal scale, See Table S27 for a list of all the questions). In Experiments 1, 2 and 3, participants answered the questionnaire by phone (to minimize drop-out rate, also to minimize close contact with participants due the covid pandemic). In Experiment 4, there were no pandemic-related restrictions and the patient filled the questionnaire in the experimenter’s presence, also to ensure that all questions were well understood. We measured autonoetic consciousness for each condition for each participant and the patient.

## Participants

In Experiment 1, 26 participants (7 male; mean age 23 ± 3.4 years) took part in the study, in Experiment 2, 29 participants (11 male, 3 gender-nonconforming, mean age 24 ± 3.4 years) and in Experiment 3, 27 participants (10 male, mean age 27 ± 3.5). All participants were right-handed as tested by the FLANDERS (Flinders Handedness Survey; FLANDERS, *100*) and reported no history of neurological or psychiatric disorder and no drug consumption in the 48h hours preceding the experiment. All participants were compensated for their participation and provided written informed consent following the local ethical committee (Cantonal Ethical Committee of Geneva: 2015-00092, and Vaud and Valais: 2016-02541) and the declaration of Helsinki (2013).

## Immersive virtual reality with motion tracking

### Immersive virtual reality

The VR paradigm and the visual stimuli were inspired and adapted from former works on EM research using immersive VR (*48*, *59*, *60*). Particular care was given to the progressive VR immersion procedure to build a strong experience of presence in the virtual environments (*101*, *102*) and to maintain it throughout the experiment. Because the immersion in VR of participant lying in an MRI scanner is particularly challenging, we based our approach on the work of (*61*), including the methods of familiarization with the virtual environment, and embodiment into a virtual body representation (*103*).

To improve the reproducibility of our paradigm, all instructions were fully automatized and provided by audio recordings through headphones. We ensured that instructions were both heard and understood during the familiarization session.

### VR display

Participants were visually immersed in VR using either a head-mounted display (Oculus Rifts S, refreshing rate 80Hz, resolution 1280 x 1440 per eye, 660 ppi; Experiments 1, 3 and 4) or MRI-compatible goggles composed of two full-HD resolution displays (1920 x 1200, 16:10 WUXGA) allowing stereoscopic rendering at 60 Hz with a diagonal field of view of 60° (Experiment 2; Visual System HD, NordicNeuroLab, Bergen, Norway).

### Motion Tracking

In Experiments 1 and 3, participants were lying down in a mock Magnetic Resonance (MR) scanner and holding custom response devices in their hands. The custom-made response device consisted of two hand-held tennis balls with integrated buttons and reflective 6-degree-of-freedom motion trackers to simultaneously maintain stable and avatar-consistent hand postures, track upper limb motion, and record participants’ answers during the in-scanner questions. Participants were wearing gloves with Velcro tape ensuring a static position of the response devices to the hand to avoid any issue related to tracker rotation and its related visual rendering in the virtual environment. We used three motion tracking cameras (Qualisys Oqus 500+m cameras with 180 Hz, 4 MegaPixel resolution) to track the devices.

In Experiment 2, participants were lying down in the MR scanner. We used a similar setup as described in (*61*). To summarize, the same tracking system is used but it is provided by six motion-tracking cameras attached to the ceiling of the MRI room to avoid any movement of the camera during the experiment and optical artifacts.

In Experiment 4, the patient was sitting on a chair, with her legs resting on a second chair in front of her, to keep the position and field of view of the scenes as similar as possible compared to the healthy participants tested in Experiments 1,2 and 3. The patient was wearing the same custom-made device used in healthy participants (Experiments 1,2,3) with the same hand position. However, because we tested the patient in another location where the motion tracking system described previously was not available, we used LEAP motion (2 cameras and 3 infrared LEDs) to track the patient’s movement.

### Software

Immersion paradigms and experimental procedures were implemented using ExVR (ExVR; Lance Florian (2019), GitHub repository: https://github.com/BlankeLab/ExVR). ExVR is a solution for designing and executing VR experiments that uses the Unity 3D engine (https://unity.com/) to perform visual rendering with realistic lighting and shading. Following principles similar to Psychopy (https://www.psychopy.org/), ExVR graphical interface allows creating complex scenes, controlling experimental variables with complex randomizations, generating logs and exporting result data compatible with standard analysis software. Additionally, for Experiment 2, ExVR also enabled the synchronization of MRI acquisition with VR experimental data.

## MRI acquisition

MR images were acquired using a 3T MRI scanner (MAGNETOM PRISMA; Siemens) using a 64-channel head coil at Campus Biotech Geneva. Each participant underwent a 5 min anatomical imaging using a T1-weighted MPRAGE sequence (TR = 2300 ms, TE = 2.25 ms, TI = 900 ms, Slice thickness = 1 mm, In-plane resolution = 1 mm × 1 mm, Number of slices = 208, FoV = 256 mm, Flip angle = 8). Encoding, BSC assessment, and recognition sessions were acquired with a whole-brain T2*-weighted Echo Planar Imaging (EPI) sequence (TR = 1500 ms, TE = 30 ms, 69 slices, flip-angle = 50°, Slice thickness = 2 mm, In-plane resolution = 2 mm × 2 mm, Multiband factor = 2, slice acquisition order = interleaved). B0 field maps were acquired during both the first and the second acquisition to correct EPI distortion due to magnetic field inhomogeneity.

### MRI preprocessing

MRI data were preprocessed with SPM12 v7487 (http://www.fil.ion.ucl.ac.uk/spm). Voxel displacement maps for the first and second sessions were calculated for each subject using pre-subtracted Phase and Magnitude Images (Short and Long echo times = 4.92 and 7.38 ms respectively, Blip direction = -1, total EPI readout time = 34.72 ms) using standard parameters. Functional images were then realigned to the first image of each session and unwarped using the voxel displacement maps with standard parameters. Images were then slice-timed to correct for time delay due to volume acquisition time using slice acquisition times recovered from DICOM raw images. Anatomical images were segmented using the unified segmentation approach (Ashburner & Friston, 2005). Functional images were corrected for bias field and then coregistered with bias field-corrected segmented anatomy using normalized mutual information. Finally, coregistered functional images were normalized using the normalization parameters estimate during unified segmentation of the anatomical images.

For the univariate GLMs specification, the functional images were smoothed using a FWHM Gaussian smoothing kernel of 5 mm. For the representational similarity analysis, we used a 2 mm-Kernel to balance spatial pattern information preservation and noise reduction (*104*).

### Categorical and trial-by-trial (TBT) General Linear Models (GLM)

Multivariate and univariate analyses used contrast maps based on subject-level random effect GLMs specified and estimated using SPM12. GLMs were estimated based on a design matrix that covered four fMRI sessions (scene encoding, BSC assessment, and two runs of scene recognition) and included boxcar regressors convolved with a canonical hemodynamic response function.

For the categorical GLM, the regressors covered the conditions of interest, nuisance covariates, and sessions. The conditions of interest for the encoding run were the 3 encoding conditions (4 × 30 s each): ENC-SYNCH1PP, ENC-ASYNCH1PP, and ENC-ASYNCH3PP, the inter-trial interval baselines (ENC-BASE, 12 × 5 s), the familiarization runs (4 × 15 s), the upper limb movement familiarization runs (3 × 30 s) for each of the 3 conditions, the in-scanner question familiarization runs (self-paced), and the button presses (0 s). The conditions of interest for BSC assessments were the 3 runs (2 × 30 s each) similar to encoding in an independent scene: BSC-SYNCH1PP, BSC-ASYNCH1PP, and BSC-ASYNCH3PP, the knife event for each condition (3 s each), the BSC questions (5 questions repeated twice, self-paced), one run (30 s) of upper limb movement without any visual stimuli, the inter-trial interval baselines (ENC_BASE, 12 × 5 s), and the button presses. For each of the recognition runs, we modeled each combination of condition, success/failure and stimulus presentation (original/changed scene) as separate regressors (for instance, REC_SYNCH1PP_Change_Success would be one possible regressor) leading to 12 possible regressors (10 s each). Attentional trials were modeled as separate regressors following the same logic (for instance, REC_SYNCH1PP_Change_Success_Catch would be one possible regressor) leading to 4 possible regressors (self-paced). The questions were modeled the same way as separate regressors. In total, for each recognition run, 32 possible regressors were modeled. Finally, similarly to other runs, we modeled inter-trial interval baselines (REC_BASE, 135 × 3 s) and button presses. The nuisance covariates were rigid translation of the head in x, y, and z direction as well as roll, pitch and yaw rotation. Finally, we added a regressor modeling individual frames exceeding 0.5 mm of framewise displacement (*105*) to regress out frames altered by excessive head movement.

We removed two participants from our MRI analysis because of excessive head movement (more than 15% of the volumes exceeding a threshold of 0.5 mm framewise displacement).

For the TBT GLM, we used the same regressors for encoding and BSC assessment as the categorical GLM but modeled each recognition trial as a single regressor to fit better statistical models of the activity and to account for possible neural repetition suppression effects.

Participant-wise GLMs were estimated for all voxels inside a common gray matter mask (SPM gray matter tissue probability map exceeding a 0.25 threshold). A high-pass filter (128 s cutoff) was applied to remove slow drifts unrelated to the paradigm.

### BSC univariate analyses

To quantify the impact of BSC manipulation on brain activity, we used a two-level analysis scheme in SPM12. From first-level contrast maps, we built a second-level within-subject ANOVA model including the three BSC conditions and derived the group-level contrasts: 2*SYNCH1PP – (ASYNCH1PP+ASYNCH3PP) to identify brain regions that were significantly activated by visuomotor and perspectival congruency compared to the other conditions. For whole-brain exploration of the effects, we used a cluster-defining threshold of p <0.001 uncorrected combined with (1) a False Discovery Rate (FDR) cluster-level correction with a threshold of p <0.05 to account for multiple comparisons. We then parcellated the cluster onto three different ROIs using the automated anatomical labeling atlas (aal *106*).

### Encoding-recognition representational similarity analysis (RSA)

To identify the brain regions displaying similar modulations of brain activity with respect to the conditions during the scene encoding and scene recognition sessions, we performed a searchlight representational similarity analysis (RSA) in a liberal gray matter mask (voxel with more than 25% of being gray matter based on SPM tissue probability maps, within an 8mm sphere (RSA;(*107*, *108*). For each participant, each session, and each voxel, we created brain representational dissimilarity matrices (bRDM) by computing the Euclidean distance between these conditions (**Fig. 3A**). We obtained a 3 × 3 matrix with 0 in the diagonal, corresponding to the distance of the condition with itself (i.e., ENC-SYNCH1PP compared to ENC-SYNCH1PP), and with the Euclidean distance of the conditions in the rest of the matrix (i.e., ENC-SYNCH1PP compared to ENC-ASYNCH1PP; **Fig. 3A**). We then computed the similarity (Z-Fisher transform of Spearman rank correlation) between the encoding bRDM and recognition bRDM for each voxel, providing an RSA brain map for each participant. Finally, we performed a permutation test by shuffling the similarity score of participants to obtain a normal distribution and select the voxel displaying significant neural similarity at a threshold of p <0.05 with a cluster size bigger than 500 voxels (**Fig. 3B**).

### Encoding-recognition similarity analysis (ERS)

To investigate the level of neural patterns of reinstatement between encoding and recognition specifically to each condition, we computed the encoding-recognition similarity score in regions of interest selected bysearchlight RSA. The ERS was computed as the Pearson correlation between vectorized neural activities corresponding (1) to encoding activity and (2) to recognition activity, for each of the conditions: SYNCH1PP, ASYNCH1PP, and ASYNCH3PP (**Fig. 3C**). It is worth mentioning that this approach is distinct from the encoding-recognition RSA: while the RSA provides a second-order isomorphism between encoding and recognition condition-related pattern, i.e., a measure of the *between-conditions* neural similarity, the ERS provides a direct linear *within-condition* similarity. Thus, while we can expect that some regions display both kinds of similarity, it is not necessarily the case by design.

We first used simple GLMs contrast maps for the calculation of ERS, to have a measure of the average reinstatement per conditions during the recognition task for each of the region of interest selected by the searchlight RSA. In a second step, we used a TBT GLMs and computed a second ERS score to measure the reinstatement of specific region of interest per trial.

### ERS as predictors of recognition performance

We used (R Core Team, 2022) and R studio (RStudio, 2022) for the analysis reported below. Linear mixed models were computed using the package *lme4* (*109*) and *modelsummary* to create table from linear mixed model results and obtain the significativity threshold (p-value) associated with each dependent variable (*110*).

To better understand the effect of conditions on ERS we used a linear mixed model approach with ERS as dependent variable (continuous) explained by conditions (3 levels: SYNCH1PP, ASYNCH1PP and ASYNCH3PP) and trials (fixed effect, continuous), with participants added as random effect and SYNCH1PP as intercept condition of the model. We applied the model separately on successful and failed trials based on our priori hypothesis that the possible effect would mainly be seen for successful trials (i.e., when the memory trace is successfully reinstated).

We used a linear mixed model approach to investigate the link between recognition performance (percentage of correct answers, dependent variable) and ERS (we extracted ERS for success and failure separately but use the average between success and failure when there was no significant difference between those two). We first compared the model with and without conditions and selected the model with lowest AIC and passing a χ 2 test (alpha = 0.05) compared to the previous model. We then derived p-values for individual effects Bonferroni corrected from the selected final model with LmerTest package.

To refine further the estimation of the relationship between recognition performance and ERS, we used a mixed effect logistic regression to explain TBT recognition performance (dependent variable, binary) as a function of conditions (fixed, 3 levels: SYNCH1PP, ASYNCH1PP and ASYNCH3PP), ERS (fixed, continuous) and stimulus presented (2 levels, original or changed scene) including their interaction for each of our ROI. We first compared the model with and without stimulus conditions and selected the model with lowest AIC and passing a χ 2 test (alpha = 0.05) compared to the previous model. We then derived p-values for individual effects from the selected final model with LmerTest package for which we added participants as a random effect and used SYNCH1PP condition as the intercept of the model.

Finally, to investigate the link between hippocampal ERS and neocortical ERS in regions indexing agency manipulation, we applied a linear mixed model with hippocampal ERS (dependent variable) as a function of conditions (fixed effect, 3 levels: SYNCH1PP, ASYNCH1PP and ASYNCH3PP), neocortical ERS (fixed, continuous) and trial number (fixed, continuous) to model habituation effects. We estimated one linear mixed model for each neocortical ERS, we then derived p-values for individual effects from the selected final model with LmerTest package for which we added participants as a random effect and used SYNCH1PP condition as the intercept of the model. We corrected the interaction results for the number of regions using a Bonferroni procedure.

## Statistical analysis

### BSC and recognition performance

Recognition performance was analyzed separately for healthy participants performing Experiments 1 and 2 (incidental encoding) and for healthy participants performing Experiment 3 (intentional encoding). Behavorial analysis was applied using R (R Core Team, 2022) and R studio (RStudio, 2022). Linear mixed models were computed using the package *lme4* (*109*)and *modelsummary* to create table from linear mixed model results (*110*).

### BSC ratings

To verify that our experimental manipulation (during encoding) impacted SoA, we used a linear mixed model with SoA ratings as dependent variable (continuous) explained by the conditions as a factor with three levels (SYNCH1PP, ASYNCH1PP, ASYNCH3PP) and participant as a random effect. The SYNCH1PP condition was used as the intercept condition for all our models. For the analysis of the incidental encoding, we added the experiment as a fixed factor (two levels: Experiment 1 and Experiment 2) to ensure there was no main difference of results between experiments. As the BSC assessment for each condition was repeated twice, we used the average rating. We report the main results in the text and all detailed tables for our mixed model can be seen in the Supplementary Materials (Table S1-S4 and S7-S10). We used the same linear mixed modeling to investigate the effect of our experimental manipulation on body ownership, control, and threat ratings. For the control questions, we averaged the ratings of the two control questions together. For the threat item, we used only the first round of ratings as dependent variable as the habituation effect was strong and is well referenced in literature (*111*, *112*).

We removed 3 participants from Experiment 1 and 2 participants from Experiment 2 due to tracking issues during BSC assessment or high ratings in the control questions which were explained by a misunderstanding of the questions by participants as revealed during post experiments feedback at the end of the recognition session. We also removed two participants from Experiment 3 because of technical issues during the experiment. Those participants were therefore not included in our BSC analysis as well as in the recognition performance analysis. We thus included a total of 76 participants across the three experiments for our BSC assessment analysis (24 Experiment 1, 27 Experiment 2, 25 Experiment 3).

### Recognition performance

To investigate if the experimental manipulation during encoding led to different levels of performance during the recognition task, we applied a mixed effect logistic regression with the binary performance score of participants as the dependent variable explained by conditions as fixed factor with three levels (SYNCH1PP, ASYNCH1PP, ASYNCH3PP) as well as the scene as a factor with three levels (ENV1, ENV2, ENV3). We added participants as a random effect and used the SYNCH1PP condition as reference condition. We used Experiment 1 as the intercept condition for our model of incidental encoding instruction and added the experiment as a factor of two levels (Experiment 1 and Experiment 2) to ensure there was no main effect of experiments. We report the main results in the text and all detailed tables for our mixed model can be seen in the Supplementary text, table section (Table S5-S6 and S11-S12). For a better visualization of the recognition performance on the figures, we plotted the recognition performance as the percent of correct answer given in the task per conditions.

To better understand whether the right upper limb movement of the participants had an effect on participant’s performance, we investigated the separated performance of participants when the objects change was on the left, versus right side of the virtual avatar. We applied a linear mixed model to explain the recognition performance (binary), with the conditions as a fixed factor with three levels (SYNCH1PP, ASYNCH1PP, ASYNCH3PP) and an interaction with the object side as a factor with two level (LEFT,RIGHT). We added the scene as a factor with three levels (ENV1,ENV2,ENV3) and the participants as a random factor. We used SYNCH1PP as the reference condition.

We removed two participants from Experiment 1, two for Experiment 2 and one for Experiment 3 because they had performance below 50% or always answered yes during the recognition task. We thus included a total of 72 participants across the three experiments for our analysis on recognition performance (22 Experiment 1, 26 Experiment 2, 24 Experiment 3).

## Patient data (Experiment 4)

In Experiment 4, we tested a 62 years old female patient (french speaker, right-handed) suffering from moderate to severe retrograde amnesia and moderate anterograde amnesia following fungal brain infection.The patient had specific deficit in autobiographical memory, more specifically in the episodic content, as tested by neuropsychologists from the hospital. The patient had bilateral lesions in the hippocampus and adjacent regions (amygdala, parahippocampus, see **Fig. 5**). The patient suffered from severe epileptic crises, which were reduced in intensity and frequency after a few months. She provided written inform consent following the local ethical committee (Cantonal Ethical Committee of Geneva: 2015-00092, and Vaud and Valais: 2016-02541) and the declaration of Helsinki (2013). An extended description of the patient’s neurological state is described in the results section and in the supplementary text.

## MRI acquisition

At the date of hospitalization, MR images were acquired using a 3T MRI scanner (MAGNETOM PRISMA; Siemens) using a 64-channel head coil at Campus Biotech Geneva. The patient underwent a 5 min anatomical imaging using a T1-weighted MPRAGE sequence (TR = 2360 ms, TE = 2.85 ms, TI = 1190 ms, Slice thickness = 1 mm, In-plane resolution = 1 mm × 1 mm, Number of slices = 200, Flip angle = 8). Prior to the scan, the patient received an gadolinium injection for better visualization of inflamed brain tissue.

Eight month after the patient’s hopsitalisation, MR images were acquired using a 3T MRI scanner (MAGNETOM PRISMA; Siemens) using a 64-channel head coil in Sion at Valais Hospital. The patient underwent the exact same T1-weighted MPRAGE sequence than the healthy participants (see MRI acquisition for the healthy participants section above).

## Volumetric analysis

To identify the regions of the medial temporal lobe affected by the infection and atrophied, we conducted a volumetry analysis using volBrain (http://volbrain.upv.es) a web-based online tool which provide information about volume of different brain regions compared to a database of around 600 healthy individual (for more information see Manjón & Coupé, 2016). We provided volBrain with the structural scan of the patient in native space. We first compared the volume of the different subpart of the hippocampus (cornus ammoni (CA) 1 to 4, the dentate gyrus, subiculum and the stratum (lacunosum SL; radiatum SR; and moleculare SM) using the HIPS pipeline of the web-based software. VolBrain preprocessed, registered the image to the MNI space and segmented the brain into different as decsribed in (*113*). As the medical report at the time of diagnosis also mentioned inflammation in the parahippocampus and amygdala, we used the standard pipeline (volbrain) of the software to investigate whether these regions were also atrophied due to the infection. For both pipeline, volBrain provide a report in pdf format describing the global volume of different tissue (white matter, grey matter, cerebrospinal fluid and intracranial volume), and more detailed measure of volume for specific brain regions (hippocampus, amygdala, parahippocampus, thalamus, putamen, caudate, see Manjón & Coupé, 2016 for the complete list). Tissue volumes are reported in cubic centimeters and as the percentage of the ratio of the structures volume with the intracranial volume (considered as 100%). Additionally, the report provides the asymmetry index, computed as the difference of volume for each region between right and left hemisphere. Normality bounds are provided by the software using their 600 healthy subjects, therefore allowing us to identify whether the amygdala, the parahippocampus, entorhinal cortex, the hippocampus and its different subpart were atrophied compared to a healthy population.

## Statistical analysis

For data from Experiment 4, we used a Crawford test to compare the patient’s BSC ratings between the three experimental conditions the ratings of healthy participants and applied the test for the three comparisons (SYNCH1PP-ASYNCH1PP, SYNCH1PP-ASYNCH3PP and ASYNCH1PP-ASYNCH3PP). We carried out the same analysis to compare the difference of recognition performance between conditions of the patient with the recognition performance of healthy participants (Crawford test applied on the three comparison SYNCH1PP-ASYNCH1PP, SYNCH1PP-ASYNCH3PP and ASYNCH1PP-ASYNCH3PP). Although the patient had an incidental encoding similar to Experiment 1 and 2, we compared her recognition performance with healthy participants who encoded the task under intentional encoding instruction (Experiment 3), because she was aware of her deficit and knew she would be tested on memory.

### Autonoetic consciousness

We first reversed the scale of four questions extracted from the EAMI questionnaire (Q1: “ How often would you estimate you have thought about this memory since it first occurred”; Q2: “ How often would you estimate you have spoken about this memory since it first occurred? “; Q3: “When you recall this event how would you describe it in terms of vividness? This can apply to the richness of sights, sounds, smells, tastes, touch, and any movements you may have made.”, Q4: “When you picture this event do you visualize it as a continuous video that plays with break, moving video clips with some breaks, one moving image or is it more like a set of snapshot with no movement, or something else?”) to have higher ratings corresponding to stronger recollection (the original EAMI questionnaire associate the lowest ratings, 1, as strong vividness and 7 a slow vividness for example). Original scaling is depicted in black in Table S27, and reversed scaling in green.

To quantify the difference of autonoetic consciousness between the patient and the young healthy participants from Experiment 3, we computed an autonoetic consciousness score for each participant and each condition, by summing the normalized ratings of the questionnaire together, to obtain one score per participant per condition. We then applied a Crawford test between the average autonoetic consciousness score of the patient between conditions, with the average autonoetic consciousness score across conditions of participants from Experiment 3.

## Acknowledgments

We would like to thank Loan Mattera and Roberto Martuzzi working at the MRI facility of Campus Biotech for their technical help during the data acquisition and the choice of the MRI sequences, and Emanuela De Falco for the insightful scientific discussion along the studies.

## Authors contributions

Conceptualization: NHM, BG, JP, FL, OB

Methodology: NHM, BG, JP, BH, MBR, OB

Investigation: NHM, BG, EF, JB, FE, MML,VA

Supervision: BG, MBR, SS, OB

Writing—original draft: NHM, BG, OB

Writing—review & editing: NHM, BG, SS, MBR, OB

## Data availability

The behavioral data will be accessible through [Git repository](https://github.com/nhmeyer/Embodied_MemoryThrough_PremotorHippocampal_Coupling) at the conclusion of the review process. While fMRI data will be shared upon request, it will not be hosted in an online repository due to privacy agreements concerning participants’ data.

