## Supplementary Materials for "Embodiment in episodic memory through premotor-hippocampal coupling"

Meyer & Gauthier *et al*

**This PDF file includes:**

Supplementary Text
Figs. S1 to S4
Tables S1 to S33
Movies S1

**Other Supplementary Materials for this manuscript include the following:**

Movies S1

**Supplementary Text**

Methods

*Experimental Design: Familiarization*

We performed a familiarization in two steps. First, immediately after entering the mock scanner (Experiment 1 and 3) or the MR scanner (Experiment 2), we made sure participants could hear the instructions given in the headset (Experiment 1 and 3) and the headphone (Experiment2). We also asked participants to perform the arm movement while displaying an outdoor empty scene and gave feedback in case the movement were too fast. This part was important for participants to get used to the movement of the arm inside the MRI and make sure they would not touch the MRI boundaries with their arm during the experiment.

Second, prior to the encoding session, participants were immersed in the four scenes (three encoding scenes and the BSC scene) but emptied from all the objects being part of the later recognition task. They were instructed to move their hands and observe the scene for 15 seconds after which they were asked one binary question “2 plus 2 equal 4” where they had to answer if this statement was correct or not, followed by a second question “How confident are you about your answer ?” to train for the two types of questions that would be asked during the experiment. Finally, they were asked to move their right arm for 30 seconds to train them for the rest of the experiment. During this time, when necessary, we interacted with the participant to tell them if the movement was too fast or not having an amplitude big enough. We used participant’s mother tongue to give the instruction when it was possible (French and English, French translations are given in supp mat) otherwise we used English. Each participant started the familiarization in the BSC scenes in the SYNCH1PP condition. The familiarization of the encoding scenes was performed in the same conditions as the one they would encode during the experiment.

*Amnestic patient with bilateral hippocampal damage is impaired in recognizing objects encoded with visuomotor and perspectival congruency*

The patient sat on a chair, with her legs resting on a second chair in front of her, approximating as much as possible the position and field of view of the scenes as tested in healthy participants ( Experiments 1-3). We used the same VR setup and head-mounted display. Another motion tracking system was used (LEAP motion; Leap Motion Controller®) because the patient comfort was tested at a hospital closer to her home.

*Patient supplementary clinical information*

End of March 2022, the patient returned home. She indicated that she initially did not recognize her apartment, where she had lived with her partner for almost one year (patient is divorced, has two daughters, and lives for 11 years together with her partner). She also noted topographical orientation deficits, with difficulties navigating in familiar environments. Concerning her autobiographical EM, she indicates that she now better remembers her work, but that she still does not remember many significant family events (i.e., births, marriage, vacations, deaths). She states that she recognizes familiar people, but does not know their name or the relationship she has with them. She is aware that she often forgets many aspects of recent discussions, because her close friends tell her that she keeps repeating the same questions.

The patient enrolled into ambulatory neuropsychological EM rehabilitation (1x/week; April to June 2022) with the aim to establish a life axis, aiming to rebuild her autobiographical memory, using photos and anecdotes, provided by the patient, her partner, one of her daughters, and two close friends. However, during these follow up sessions and interviews the patient's EM remains severely impaired; she is not able to produce autobiographical events and only produces a few semantical events (which remained deficient). After three months of rehabilitation, the patient starts to orient her life axis and can provide certain elements, but is not able to evoke or relive any of these life events. Indications by others do not help her to relive these life events either. She uses a life booklet, in which she noted with her family and friends most of her key life events.

### Results

***Incidental encoding. Visuomotor and perspectival congruency for incidental encoding does not modulate object recognition (behavior, Experiment 1 and 2):*** We also compared the effect of recognition performance between scenes to ensure that each scene had the same level of difficulty. We found that scene 2 was significantly easier compared to the two other scenes in incidental encoding , Experiment 1 and 2; estimate = 0.22, z = 3.33, p = 0.001) but not in intentional encoding (Experiment 3 ; estimate = 0.013, z = 0.12, p = 0.9). Therefore, we included the scene as fixed effect in our analysis to take this bias into account. However, since the association between condition and scene was pseudorandomized between participants this effect should not affect our findings.

#### *ERS analysis. Reinstatement in the left hippocampus is higher for visuomotor and perspectival congruency and indexes recognition memory.*

*Model selection to explain recognition performance with hippocampal ERS*

To better understand the link between hippocampal ERS and memory, we compared a model which explains recognition performance using hippocampal ERS and conditions (Model 1) with a model considering only hippocampal ERS irrespective of conditions (Model 0). We found that both models were equally good (i.e. had the same AIC; m1 AIC = -159.83, m0 AIC = -159.73, $X_2$= 8.10, p = 0.088, Table S28). Therefore we used the model with the smaller number of parameters (Model 0) for further analysis.

*Hippocampal ERS and performance, Trial-by-Trial*

The positive correlation between left hippocampal ERS and recognition performance was found for average hippocampal ERS (per session average of successful and failed trials) and for the overall recognition performance (percent of correct answers). To investigate whether this relation holds for single trials, we applied a logistic mixed effect model to investigate trial-by-trial recognition performance with trial-by-trial hippocampal ERS (as described in the main text). This analysis revealed a significant triple interaction between condition, stimulus (same scene than the one at encoding versus changed scene), and left hippocampal ERS, when SYNCH1PP was compared to ASYNCH3PP (estimate = -2.04, z = -2.6, p = 0.009, Table S29-31). P*ost-hoc* analysis revealed that the significant effect was driven by the significantly positive relationship between recognition performance and left hippocampal ERS in SYNCH1PP (**Fig. 4B,** Table S32), but only when the stimulus presented was the same scene as the one observed at encoding (estimate = 1.7, z = 3.5, p <0.001). The relation between hippocampal ERS and recognition performance was not significant for the ASYNCH3PP conditions (Table S33). This shows that only the main experimental condition with visuomotor and perspectival congruency, associated activity in left hippocampus with recognition performance on a trial by trial basis.

***Hippocampal-neocortical interactions revealed by ERS are modulated by visuomotor and perspectival congruency***

We found that participant's SoA was correlated with the activity of the BSC regions at encoding (Premotor left: r = 0.22, df = 79, t = 1.98, p = 0.05, right SMA: r = 0.35, df = 79, t = 3.38, p = 0.001, left SMA: r = 0.3, df = 79, t = 2.81, p = 0.006 , suggesting that these regions identified using the contrast (SYNCH1PP > ASYNCH1PP +ASYNCH3PP) are involved in the subjective outcome of the BSC manipulation (SoA).

*Amnestic patient with bilateral hippocampal damage is impaired in recognizing objects encoded with visuomotor and perspectival congruency*

We tested the patient five months after her hospitalization. At that time, The patient's scored 25 at the Montreal Cognitive Assessment test, with 100% correct answer on the memory part of the test although she still suffered from autobiographical memory deficit.

The difference in the patient's SoA ratings between the SYNCH1PP condition and both ASYNCH1PP and ASYNCH3PP was modulated in the same way as observed in the healthy participants and her sensitivity to the manipulation was even higher compared to healthy participants as tested using Crawford test, due to larger SoA differences across conditions (SYNCH1PP compared to ASYNCH1PP: mean = 0.05, sd ± = 0.16, p < .001; ASYNCH1PP compared to ASYNCH3PP: mean = 0.03, sd ± = 0.16, p = 0.004). The SoA difference between SYNCH1PP and ASYNCH3PP was not significantly different compared to healthy participants but going in the same direction (SYNCH1PP-ASYNCH3PP: mean = 0.08, sd ± = 0.18, p = 0.134). Similar findings were obtained for ownership ratings (SYNCH1PP-ASYNCH1PP: mean = 4.00e-03, sd ± = 0.21, p = 0.002). The other difference between conditions were in the same range than the healthy participants rating (SYNCH1PP-ASYNCH3PP: mean = 0.12, sd ± = 0.23, p < .001; ASYNCH1PP-ASYNCH3PP: mean = 0.12, sd ± = 0.21, p = 0.314). The patient's ratings for control items were low, did not differ between conditions, and also did not differ from those of healthy participants (SYNCH1PP-ASYNCH1PP: mean = -7.41e-03, sd ± = 0.09, p = 0.257;

SYNCH1PP-ASYNCH3PP: mean = 0.02, sd ± = 0.07, $p = 0.295$; ASYNCH1PP-ASYNCH3PP: mean = 0.03, sd ± = 0.11, $p = 0.438$).

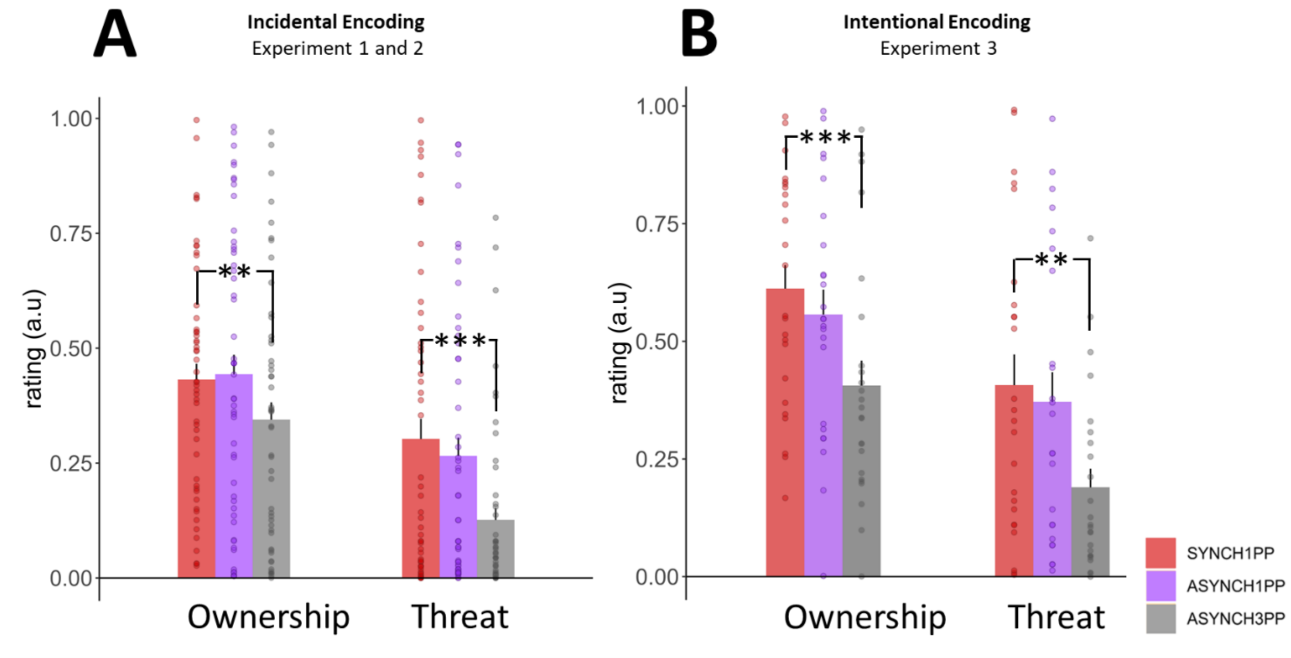


**Fig. S1.**

**BSC ratings during encoding of scenes under different visuomotor conditions**. (**A**) Participants had a higher body ownership rating and were more afraid of the under visuomotor and perspectival congruency (SYNCH1PP, red) compared to visuomotor and perspectival mismatch (ASYNCH3PP, grey), no difference was observed between visuomotor and perspectival congruency (SYNCH1PP) and visuomotor mismatch (ASYNCH1PP, purple). **,*** indicates significance level with p-value <0.01, < 0.001 respectively, as tested with a linear mixed model ; N = 50. (**B**) Participants had a higher body ownership and were more afraid of the threat under visuomotor and perspectival congruency (SYNCH1PP, red) compared to visuomotor and perspectival mismatch (ASYNCH3PP, grey), no difference was observed between visuomotor and perspectival congruency (SYNCH1PP) and visuomotor mismatch (ASYNCH1PP, purple). **, *** indicates significance level with p-value <0.01 , < 0.001 respectively as tested with a linear mixed model N = 25.

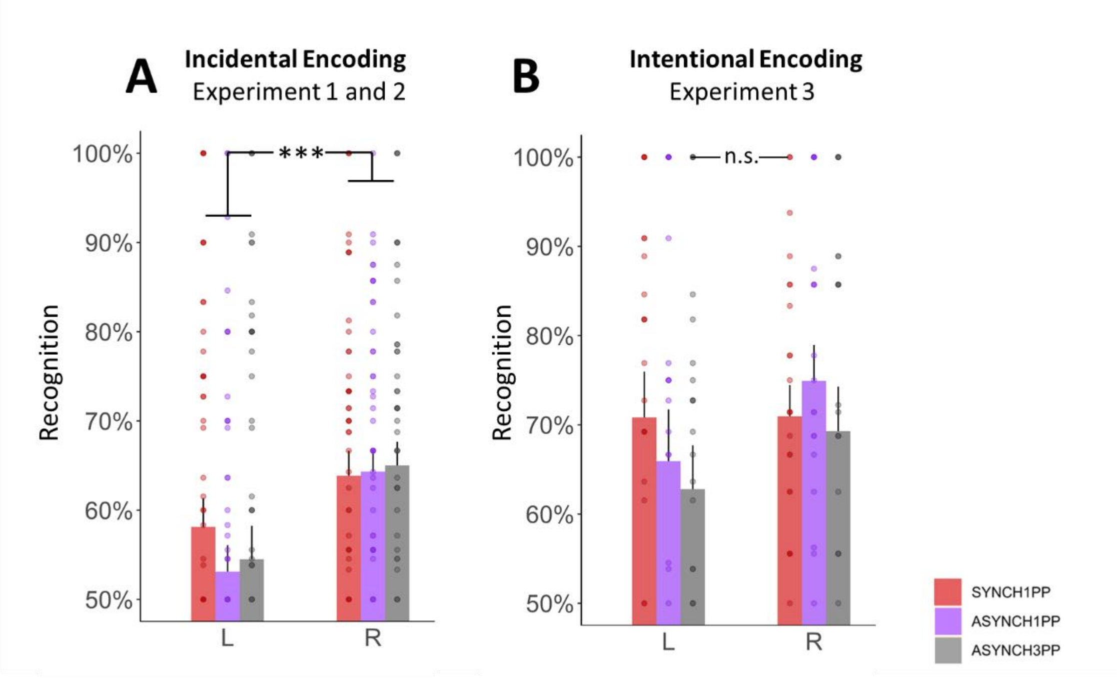


**Fig. S2.**

**Effect of objects changes laterality on recognition**. (**A**) There was a significant main effect of object side but no interaction between conditions and object side under the incidental encoding instruction (experiment 1& 2) as tested with linear mixed model, N = 50. *** indicates significance level with p-value <0.001. (**B**) There was no effect of object laterality under intentional encoding instruction (experiment 3). linear mixed model, N = 25.

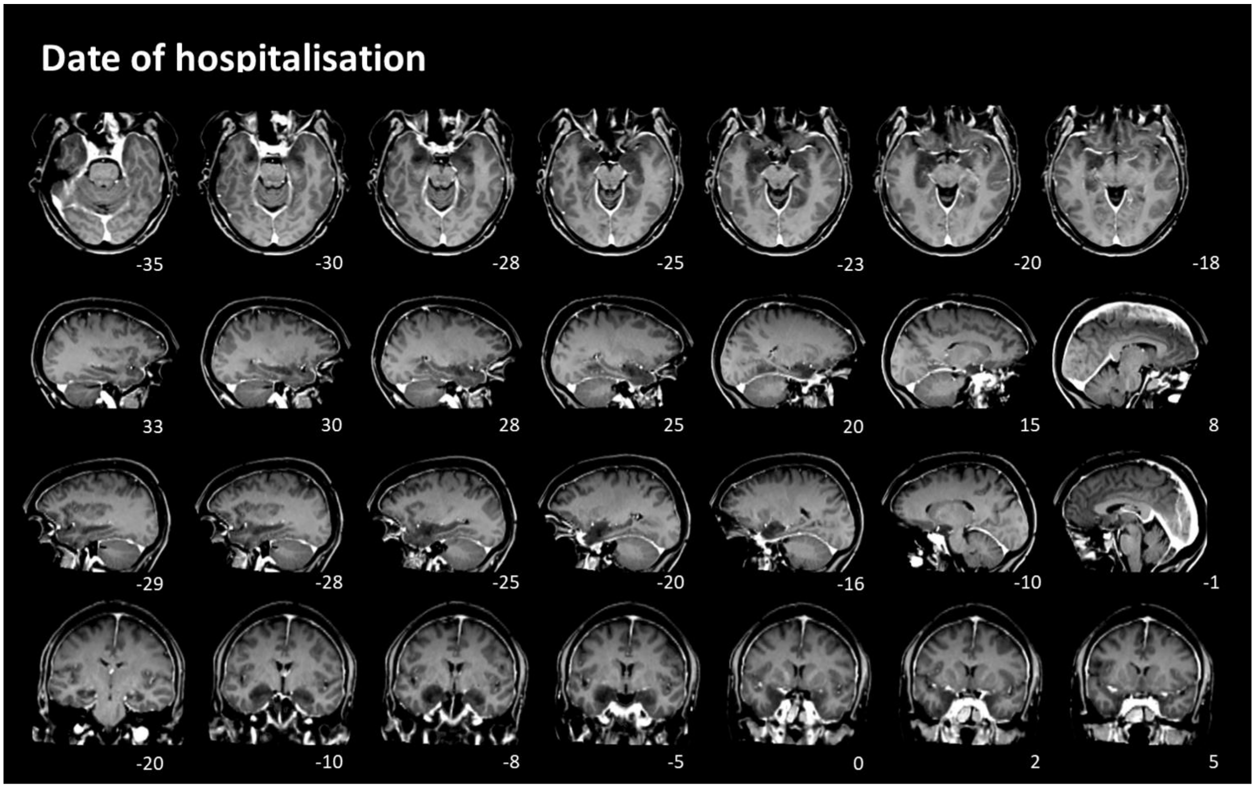


**Fig. S3.**

**Patient’s lesion on the day of hospitalisation.** Anatomical scan of the patient on the day of the hospitatlisaton acquired with a Siemens MR-scanner (3T). dark regions around the left hippocampus show sign of inflammation.

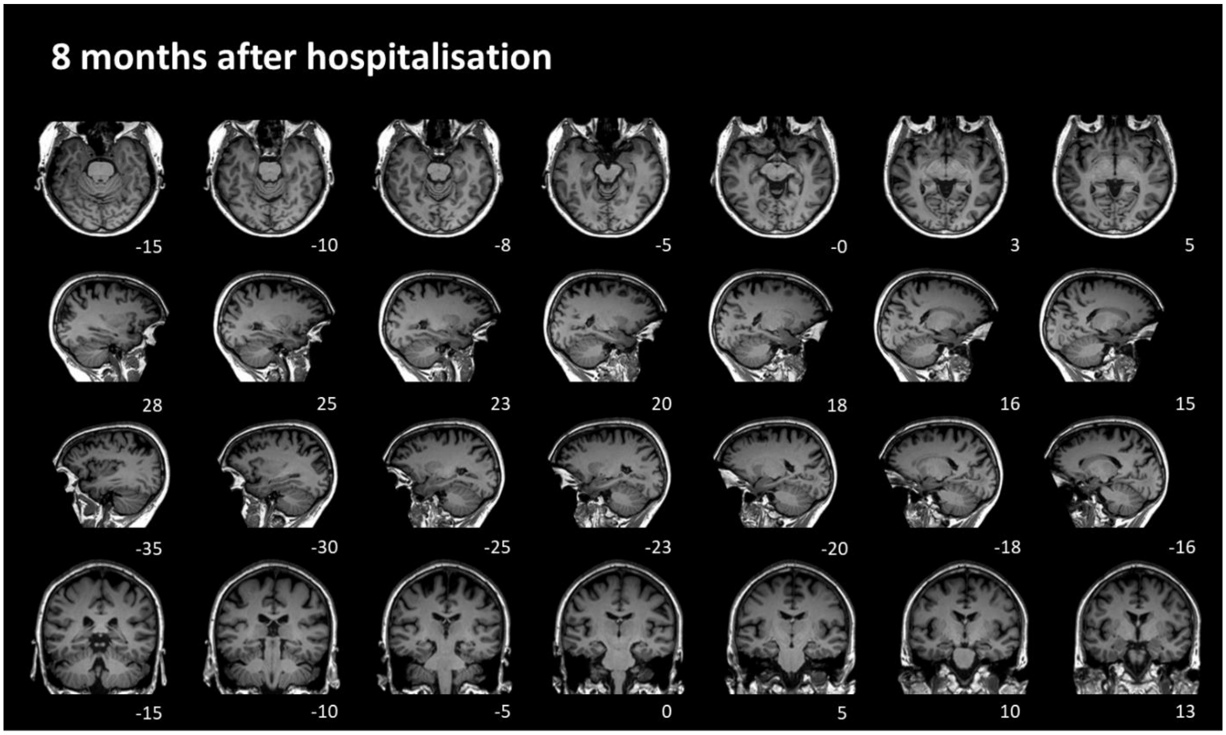


**Fig. S4.**

**Patient's lesion eight month after hospitalisation.** Anatomical scan of the patient taken eight month after the diagnosis. Clear amelioration of the inflammation around the hippocampal regions based on clinical report, although the patient did not recover from her amnestic deficit.

| | Estimate | t-value | p-value |
|---|---|---|---|
| **(Intercept)** | 0.654 | 12.383 | <0.001 *** |
| **factor(Conditions)ASYNCH1PP** | -0.067 | -2.931 | 0.003** |
| **factor(Conditions)ASYNCH3PP** | -0.065 | -2.841 | 0.005** |
| **factor(XP)MRI** | -0.045 | -0.652 | 0.515 |

**Table S1.**

**Effect of conditions on sense of agency ratings in Experiment 1 and 2:** Agency ~ Conditions + random(Participants).

| | Estimate | t-value | p-value |
| --- | --- | --- | --- |
| **(Intercept)** | 0.464 | 8.285 | <0.001 *** |
| **factor(Conditions)ASYNCH1PP** | 0.012 | 0.387 | 0.699 |
| **factor(Conditions)ASYNCH3PP** | -0.087 | -2.903 | 0.004 ** |
| **factor(XP)MRI** | -0.833 | -0.833 | 0.405 |

**Table S2.**

**Effect of conditions on ownership ratings in Experiment 1 and 2:** Ownership ~ Conditions + random(Participants).

| | Estimate | t-value | p-value |
|---|---|---|---|
| **(Intercept)** | 0.289 | 5.557 | <0.001 *** |
| **factor(Conditions)ASYNCH1PP** | -0.037 | -0.936 | 0.349 |
| **factor(Conditions)ASYNCH3PP** | -0.176 | -4.451 | <0.001 *** |
| **factor(XP)MRI** | 0.024 | 0.384 | 0.701 |

**Table S3.**

**Effect of conditions on threat ratings in Experiment 1 and 2:** Threat ~ Conditions + random(Participants).

| | Estimate | t-value | p-value |
|---|---|---|---|
| **(Intercept)** | 0.068 | 2.678 | 0.007 ** |
| **factor(Conditions)ASYNCH1PP** | 0.012 | 0.942 | 0.346 |
| **factor(Conditions)ASYNCH3PP** | -0.013 | -1.043 | 0.297 |
| **factor(XP)MRI** | 0.061 | 1.829 | 0.067 |

**Table S4.**

**Effect of conditions on control ratings for experimental bias in Experiment 1 and 2:** Control ~ Conditions + random(Participants).

| | Estimate | z-value | p-value |
|---|---|---|---|
| **(Intercept)** | 0.643 | 6.925 | <0.001 *** |
| **factor(Conditions)ASYNCH1PP** | 0.025 | 0.355 | 0.723 |
| **factor(Conditions)ASYNCH3PP** | 0.064 | 0.929 | 0.353 |
| **factor(Environment)ENV2** | 0.207 | 2.969 | 0.003 ** |
| **factor(Environment)ENV3** | -0.002 | -0.031 | 0.975 |
| **factor(XP)MRI** | -0.025 | -0.243 | 0.808 |

**Table S5.**

**Effect of conditions on performance (Experiment 1-2)**: Performance ~ Conditions + Scene + random(Participants).

| | Estimate | z-value | p-value |
|---|---|---|---|
| (Intercept) | 0.212 | 1.348 | 0.178 |
| factor(Conditions)ASYNCH1PP | -0.201 | -1.421 | 0.155 |
| factor(Conditions)ASYNCH3PP | -0.177 | -1.238 | 0.216 |
| Factor(Laterality) R | 0.288 | 2.028 | 0.043 * |
| factor(Environment)ENV2 | 0.645 | 6.440 | <0.001 *** |
| factor(Environment)ENV3 | 0.091 | 0.917 | 0.359 |
| factor(XP)MRI | -0.277 | -1.946 | 0.052 |
| factor(Conditions)ASYNCH1PP × factor(Laterality)R | 0.145 | 0.737 | 0.461 |
| factor(Conditions)ASYNCH3PP × factor(Laterality)R | 0.162 | 0.820 | 0.412 |

**Table S6.**

**Effect of object laterality on performance (Experiment 1-2)**: Performance ~ Conditions*ObjectLaterality(L/R) + Scene + random(Participants).

| | Estimate | t-value | p-value |
|---|---|---|---|
| **(Intercept)** | 0.695 | 13.684 | <0.001 *** |
| **factor(Conditions)ASYNCH1PP** | -0.042 | -1.202 | 0.229 |
| **factor(Conditions)ASYNCH3PP** | -0.111 | -3.159 | 0.002 ** |

**Table S7.**

**Effect of conditions on sense of agency ratings in Experiment 3:** Agency ~ Conditions + random(Participants).

| | Estimate | t-value | p-value |
|---|---|---|---|
| **(Intercept)** | 0.612 | 11.760 | <0.001 *** |
| **factor(Conditions)ASYNCH1PP** | -0.055 | -1.226 | 0.220 |
| **factor(Conditions)ASYNCH3PP** | -0.206 | -4.590 | <0.001 *** |

**Table S8.**

**Effect of conditions on ownership ratings in Experiment 3:** Ownership ~ Conditions + random(Participants).

| | Estimate | t-value | p-value |
|---|---|---|---|
| **(Intercept)** | 0.407 | 7.136 | <0.001 *** |
| **factor(Conditions)ASYNCH1PP** | -0.035 | -0.508 | 0.612 |
| **factor(Conditions)ASYNCH3PP** | -0.217 | -3.126 | 0.002 ** |

**Table S9.**

**Effect of conditions on threat ratings in Experiment 3:** Threat ~ Conditions + random(Participants).

| | Estimate | t-value | p-value |
|---|---|---|---|
| **(Intercept)** | 0.179 | 5.600 | <0.001 *** |
| **factor(Conditions)ASYNCH1PP** | -0.010 | -0.557 | 0.578 |
| **factor(Conditions)ASYNCH3PP** | -0.028 | -1.503 | 0.133 |

**Table S10.**

**Effect of conditions on control ratings for experimental bias in Experiment 3:** Control ~ Conditions + random(Participants).

| | Estimate | z-value | p-value |
|---|---|---|---|
| **(Intercept)** | 1.211 | 8.607 | <0.001 *** |
| **factor(Conditions)ASYNCH1PP** | -0.328 | -3.058 | 0.002 ** |
| **factor(Conditions)ASYNCH3PP** | -0.316 | -2.949 | 0.003 ** |
| **factor(Environment)ENV2** | 0.013 | 0.119 | 0.905 |
| **factor(Environment)ENV3** | 0.100 | 0.943 | 0.346 |

**Table S11.**

**Effect of conditions on performance (Experiment 3)**: Performance ~ Conditions + Scene + random(Participants).

| | Estimate | z-value | p-value |
|---|---|---|---|
| **(Intercept)** | 0.671 | 3.087 | 0.002 ** |
| **factor(Conditions)ASYNCH1PP** | -0.399 | -1.87 | 0.062 |
| **factor(Conditions)ASYNCH3PP** | -0.398 | -1.864 | 0.062 |
| **Factor(Laterality) R** | 0.037 | 0.168 | 0.867 |
| **factor(Environment)ENV2** | 0.5 | 283 | 0.001 ** |
| **factor(Environment)ENV3** | 0.255 | 1.662 | 0.095 |
| **factor(Conditions)ASYNCH1PP × factor(Laterality)R** | 0.502 | 0.737 | 0.097 |
| **factor(Conditions)ASYNCH3PP × factor(Laterality)R** | 0.194 | 0.650 | 0.515 |

**Table S12.**

**Effect of object laterality on performance (Experiment 3)**: Performance ~ Conditions*ObjectLaterality(L/R) + Scene + random(Participants).

| | Estimate | t-value | p-value |
|---|---|---|---|
| **(Intercept)** | 0.039 | 1.700 | 0.089 |
| **factor(Conditions)ASYNCH1PP** | -0.045 | -2.570 | 0.010 |
| **factor(Conditions)ASYNCH3PP** | -0.045 | -2.596 | 0.009 ** |
| **factor(Environment)ENV2** | -0.012 | -0.668 | 0.504 |
| **factor(Environment)ENV3** | -0.006 | -0.323 | 0.747 |

**Table S13.**

**Effect of conditions on hippocampal ERS**: Hippocampal ERS Success ~ Conditions + Scene + random(Participants).

| | Estimate | t-value | p-value |
|---|---|---|---|
| **(Intercept)** | 0.008 | 0.327 | 0.743 |
| **factor(Conditions)ASYNCH1PP** | -0.048 | -2.769 | 0.006 ** |
| **factor(Conditions)ASYNCH3PP** | -0.011 | -0.637 | 0.524 |
| **factor(Environment)ENV2** | 0.033 | 1.910 | 0.056 |
| **factor(Environment)ENV3** | 0.001 | 0.083 | 0.934 |

**Table S14.**

**Effect of conditions on middle temporal gyrus ERS**: middle temporal gyrus ERS Success ~ Conditions + Scene + random(Participants).

| | Estimate | t-value | p-value |
|---|---|---|---|
| **(Intercept)** | 0.039 | 2.254 | 0.024 * |
| **factor(Conditions)ASYNCH1PP** | -0.010 | -0.478 | 0.633 |
| **factor(Conditions)ASYNCH3PP** | -0.033 | -1.628 | 0.103 |
| **factor(Environment)ENV2** | -0.009 | -0.462 | 0.644 |
| **factor(Environment)ENV3** | -0.002 | -0.120 | 0.904 |

**Table S15.**

**Effect of conditions on orbitofrontal ERS**: orbitofrontal ERS Success ~ Conditions + Scene + random(Participants).

| | Estimate | t-value | p-value |
|---|---|---|---|
| **(Intercept)** | 0.061 | 1.795 | 0.073 |
| **factor(Conditions)ASYNCH1PP** | -0.001 | -0.043 | 0.966 |
| **factor(Conditions)ASYNCH3PP** | -0.030 | -1.590 | 0.112 |
| **factor(Environment)ENV2** | 0.003 | 0.155 | 0.877 |
| **factor(Environment)ENV3** | -0.029 | -1.544 | 0.123 |

**Table S16.**

**Effect of conditions on calcarine ERS**: Calcarine ERS Success ~ Conditions + Scene + random(Participants).

| | *AIC* | *BIC* | *logLik* | *deviance* | *Chisq* | *Df* | *Pr(>Chisq)* |
|---|---|---|---|---|---|---|---|
| ***Model 0*** | 159.731 | 146.071 | 85.86549 | 171.731 | | | |
| ***Model 1*** | 159.834 | 137.068 | 89.91708 | 179.834 | 8.103189 | 4 | 0.087871 |

**Table S17.**

**Model comparison : performance explained by ERS Hippocampus with or without conditions.** (Model 0) Performance ~ ERS Hippocampus + Scene + random(Participants). **(Model1)** Performance ~ ERS Hippocampus * Conditions + Scene + random(Participants)

| | Estimate | t-value | p-value |
|---|---|---|---|
| **(Intercept)** | 0.661 | 40.842 | <0.001*** |
| **ERS** | 0.291 | 2.723 | 0.006 ** |
| **factor(Environment)ENV2** | 0.052 | 2.748 | 0.006 ** |
| **factor(Environment)ENV3** | -0.008 | -0.409 | 0.682 |

**Table S18.**

**Effect of hippocampal ERS on performance**: *Performance ~ Hippocampal ERS + Scene + random(Participants).*

| | [x y z] | k | p-value |
|---|---|---|---|
| **Cluster** | [6 -2 54] | 856 | 0.02 |

**Table S19.**

***Cluster sensitive to the BSC manipulation at encoding:****Synch1pp-(Asynch1PP+Asynch3PP) BSC session.*

| | Estimate | t-value | p-value |
|---|---|---|---|
| **(Intercept)** | 0.66 | 30.77 | <0.001*** |
| **ERS** | 0.08 | 0.6 | 0.55 |
| **factor(Environment)ENV2** | 0.04 | 1.8 | 0.08 |
| **factor(Environment)ENV3** | -0.02 | -0.61 | 0.54 |

**Table S20.**

**Effect of middle temporal gyrus ERS on performance**: Performance ~ Middle temporal gyrus ERS + Scene + random(Participants).

| | Estimate | t-value | p-value |
|---|---|---|---|
| **(Intercept)** | 0.019 | 0.877 | 0.380 |
| **factor(Conditions)ASYNCH1PP** | -0.013 | -1.147 | 0.252 |
| **factor(Conditions)ASYNCH3PP** | -0.008 | -0.714 | 0.475 |
| **ERS** | 0.199 | 7.404 | <0.001*** |
| **Trials** | -0.002 | -4.674 | <0.001*** |
| **factor(Conditions)ASYNCH1PP × ERS** | -0.192 | -5.264 | <0.001*** |
| **factor(Conditions)ASYNCH3PP × ERS** | -0.069 | -1.896 | 0.058 |

**Table S21.**

**Left dPMC and hippocampal ERS coupling**: *Hippocampal ERS ~ Premotor ERS * Conditions +Trials + random(Participants).*

| | Estimate | t-value | p-value |
|---|---|---|---|
| **(Intercept)** | 0.035 | 1.269 | 0.205 |
| **ERS** | 0.194 | 6.765 | <0.001*** |
| **Trials** | -0.003 | -3.730 | <0.001*** |

**Table S22.**

**Left dPMC and hippocampal ERS coupling for SYNCH1PP**: *Hippocampal ERS ~ Premotor ERS +Trials + random(Participants).*

| | Estimate | t-value | p-value |
|---|---|---|---|
| **(Intercept)** | 0.001 | 0.045 | 0.964 |
| **ERS** | -0.038 | -1.366 | 0.172 |
| **Trials** | -0.002 | -2.254 | 0.024 * |

**Table S23.**

**Left dPMC and hippocampal ERS coupling for ASYNCH1PP**: *Hippocampal ERS ~ Premotor ERS +Trials + random(Participants).*

| | Estimate | t-value | p-value |
|---|---|---|---|
| **(Intercept)** | 0.012 | 0.564 | 0.573 |
| **factor(Conditions)ASYNCH1PP** | -0.019 | -1.692 | 0.091 |
| **factor(Conditions)ASYNCH3PP** | -0.012 | 2.454 | 0.262 |
| **ERS** | 0.061 | 7.404 | 0.014 * |
| **Trials** | -0.001 | -2.961 | 0.003 ** |
| **factor(Conditions)ASYNCH1PP × ERS** | 0.007 | 0.216 | 0.829 |
| **factor(Conditions)ASYNCH3PP × ERS** | 0.006 | 0.171 | 0.864 |

**Table S24.**

**Left SMA and hippocampal ERS coupling**: Hippocampal ERS ~ Left SMA ERS *Conditions +Trials + random(Participants).

| | Estimate | t-value | p-value |
|---|---|---|---|
| **(Intercept)** | 0.020 | 0.946 | 0.344 |
| **factor(Conditions)ASYNCH1PP** | -0.019 | -1.674 | 0.094 |
| **factor(Conditions)ASYNCH3PP** | -0.012 | -1.106 | 0.269 |
| **ERS** | 0.006 | 0.260 | 0.795 |
| **Trials** | -0.002 | -3.962 | <0.001*** |
| **factor(Conditions)ASYNCH1PP × ERS** | -0.002 | -0.059 | 0.953 |
| **factor(Conditions)ASYNCH3PP × ERS** | 0.046 | 1.366 | 0.172 |

**Table S25.**

**Right SMA and hippocampal ERS coupling**: Hippocampal ERS ~ Right SMA ERS *Conditions +Trials + random(Participants).

| Volumes | Total ($cm^3$/%) | Right ($cm^3$/%) | Left ($cm^3$/%) | Assymetry (%) Right - Left |
|---|---|---|---|---|
| **Hippocampus** | 2.46 / (0.1944) [ 0.28 - 0.43] | 1.12 / (0.0883) [ 0.14 - 0.22] | 1.34 / (0.1061) [ 0.14 - 0.22] | -18.2331 [-16.78 - 10.90] |
| ***CA1*** | 0.93 / (0.0738) [ 0.10 - 0.15] | 0.44 / (0.0349) [ 0.05 - 0.08] | 0.49 / (0.0389) [ 0.05 - 0.08] | -11.0065 [-21.32 - 15.49] |
| ***CA2-CA3*** | 0.16 (0.0128) [ 0.02 - 0.03] | 0.08 / (0.0062) [ 0.01 - 0.02] | 0.08 / (0.0066) [ 0.01 - 0.02] | -5.5468 [-66.14 - 27.57] |
| ***CA4-DG*** | 0.55 / (0.0434) [ 0.07 - 0.11] | 0.24 / (0.0189) [ 0.03 - 0.06] | 0.31 / (0.0245) [ 0.03 - 0.06] | -26.0628 [-66.14 - 27.57] |
| ***SR-SL-SM*** | 0.46 / (0.0360) [ 0.03 - 0.04] | 0.20 / (0.0160) [ 0.03 - 0.04] | 0.25 / (0.0200) [ 0.03 - 0.04] | -22.5932 [-24.20 - 23.20] |
| ***Subiculum*** | 0.36 / (0.0285) [ 0.03 - 0.05] | 0.16 / (0.0124) [ 0.02 - 0.03] | 0.20 / (0.0160) [ 0.02 - 0.03] | -25.2172 [-16.68 - 29.38] |
| **Amgydala** | 1.68 / (0.135) [0.117, 0.172] | 0.89 / (0.072) [0.059, 0.086] | 0.79 / (0.063) [0.057, 0.087] | 12.5550 [-11.868, 13.681] |
| **Entorhinal area** | 3.80 / (0.305) [0.226, 0.348] | 2.06 / (0.165) [0.113, 0.178] | 1.74 / (0.140 [0.105, 0.178] | 16.8496 [-19.903, 25.976] |
| **Parahippocampal gyrus** | 5.12 / (0.411) [0.332, 0.500] | 2.44 / (0.196) [0.157, 0.248] | 2.69 / (0.216) [0.170, 0.261] | -9.7990 [-24.494, 10.192] |

**Table S26.**

**Volumetry analysis of the inflamed regions from the diagnosis date eight months after the infection.** Eight months after the infection, the patient had significantly smaller bilateral hippocampi volume but spared amygdala, entorhinal cortex and parahippocampal gyrus as compared with 600 healthy participants using the volbrain software. For each region, the software provide the absolute volume in cubic centimeters ($cm^3$) and in percent, computed as the ratio between the region's volume and the intracranial volume (considered as 100%). Number in brackets correspond to the normative value of neurologically intact population provided by the software. CA = cornu ammoni, DG = dentate gyrus, SR = stratum radiatum, SL = stratum lacunosum, SM = stratum moleculare.

| Statement | Scale | Reference |
|---|---|---|
| **My memory for this event involves sound** | 1-7 little/A lot | MCQ |
| **My memory for this event involves smell** | 1-7 little/A lot | MCQ |
| **My memory for this event involves touch** | 1-7 little/A lot | MCQ |
| **The overall tone of the memory is** | Negative/Neutral/Positive | MCQ |
| **In this event I was** | An observer /A participant | MCQ |
| **I remember the event through my own eyes as during the event** | 1-7 Not at all/Definitely | MCQ |
| **When you picture this event do you visualize it as a continuous video that plays with break, moving video clips with some breaks, one moving image or is it more like a set of snapshot with no movement, or something else?** | 1-7<br>One smooth video/video clips with breaks/one moving image/snapshot in sequence/one static snapshot/Hazy image/no image<br>no image / Hazy image/one static snapshot /snapshot in sequence /one moving image /video clips with breaks /One smooth video | EAMI |
| **How often would you estimate you have thought about this memory since it first occurred?** | 1- 4<br>Frequently/Occasionnaly/Rarely/Never<br>Never/Rarely/Occasionnaly/Frequently | EAMI |
| **How often would you estimate you have spoken about this memory since it first occurred?** | 1-4<br>Frequently/Occasionnaly/Rarely/Never<br>Never/Rarely/Occasionnaly/Frequently | EAMI |
| **When you recall this event are you viewing the scene through your « own eyes » or can you see yourself in the memory from a third-person perspective?** | Own eyes/Mixture/Third person/something different/no imagery | EAMI |
| **When you recall this event how would you describe it in terms of vividness? This can apply to the richness of sights, sounds, smells, tastes, touch, and any movements you may have made.** | 1-7 very vivid/very vague<br>1-7 very vague/very vivid | EAMI |
| **The relative spatial arrangement of people in my memory for the event is** | 1-7 Vague/Distinct | MCQ |
| **My memory for the time when the event takes place is** | 1-7 Vague/Distinct | MCQ |
| **When I remember the event, I see myself entirely in the scene as if I was watching a movie** | 1-7 Not at all/Definitely | MCQ |
| **When I think about or tell this memory, I feel like I relive it as it happened** | 1-7 Not at all/Definitely | MCQ |
| **I remember the movements and gestures I made with my body at the time of the event** | 1-7 / Vague/Distinct | In-house |
| **My memory for this event is** | 1-7 Dim/Clear | MCQ |
| **My memory for this event involves visual details** | 1-7 Little/ A lot | MCQ |
| **My memory for this event is** | 1-7 Sketchy/very detailed | MCQ |
| **My memory for the location where the event takes place is** | 1-7 Vague/Distinct | MCQ |
| **Relative spatial arrangement of objects in my memory for the event is** | 1-7 Vague/Distinct | MCQ |
| **When you think about this event now, do you re-experience any of the emotion you originally felt at the time? To what extent are you re-experiencing this emotion as a percentage?** | 0/25/50/75/100% | EAMI |
| **To what extent are you re-experiencing this memory as a percentage?** | 0/25/50/75/100% | EAMI |
| **Would you say you are reliving this memory or looking back on it?** | Reliving/Looking back | EAMI |

| | | |
|---|---|---|
| **I remember how I felt at the time when the event took place** | 1-7 Not at all/Definitely | MCQ |
| **I remember what I thought at the time** | 1-7 Not at all/Definitely | MCQ |

**Table S27.**

**Autonoetic consciousness questionnaire**. Scale from original questionnaire is indicated in black, new scale is indicated in green.

| | *AIC* | *BIC* | *logLik* | *deviance* | *Chisq* | *Df* | *Pr(>Chisq)* |
|---|---|---|---|---|---|---|---|
| ***Model 0*** | *3580.075* | *3627.8* | *1782.04* | *3564.075* | | | |
| ***Model 1*** | *3416.513* | *3500.031* | *1694.26* | *3388.513* | *175.5622* | *6* | *<0.001* |

**Table S28.**

**Model comparison: performance (trial-by-trial) explained by hippocampal ERS and conditions or hippocampal ERS, conditions, and stimulus type.** (Model 0) Performance (binomial) ~hippocampal ERS * Conditions + Trials +random(Participants).(Model 1) Performance (binomial) ~hippocampal ERS * Conditions *Stimulus+ Trials +random(Participants)

| | Estimate | z-value | p-value |
|---|---|---|---|
| (Intercept) | -0.368 | -2.910 | <0.001*** |
| factor(Conditions)ASYNCH1PP | 0.088 | 0.661 | 0.509 |
| factor(Conditions)ASYNCH3PP | 0.104 | 0.780 | 0.435 |
| ERS | -0.411 | -1.168 | 0.243 |
| Factor(Stimulus)NoChange | 1.009 | 6.874 | <0.001*** |
| Trials | 0.028 | 7.684 | 0.001 ** |
| factor(Conditions)ASYNCH1PP × ERS | 0.650 | 1.293 | 0.196 |
| factor(Conditions)ASYNCH3PP × ERS | 0.684 | 1.342 | 0.180 |
| factor(Conditions)ASYNCH1PP × Stimulus(NoChange) | 0.168 | 0.791 | 0.429 |
| factor(Conditions)ASYNCH3PP × Stimulus(NoChange) | 0.056 | 3.524 | 0.787 |
| ERS x factor(Stimulus) NoChange | 1.862 | 3.524 | <0.001*** |
| factor(Conditions)ASYNCH1PP × ERSx factor(Stimulus) NoChange | -1.073 | -1.380 | 0.168 |
| factor(Conditions)ASYNCH3PP × ERSx factor(Stimulus) NoChange | -2.043 | -2.604 | 0.009 ** |

**Table S29.**

**Effect of hippocampal ERS , conditions and type of stimuli (Original scene or changed scene) on performance:** Performance ~ Hippocampal ERS Conditions*Stim +Trials + random(Participants)

| | Estimate | z-value | p-value |
|---|---|---|---|
| **(Intercept)** | -0.105 | -0.686 | 0.493 |
| **factor(Conditions)ASYNCH1PP** | 0.089 | 0.670 | 0.503 |
| **factor(Conditions)ASYNCH3PP** | 0.112 | 0.835 | 0.403 |
| **ERS** | -0.509 | -1.429 | 0.153 |
| **Trials** | 0.028 | 7.684 | 0.001 ** |
| **factor(Conditions)ASYNCH1PP × ERS** | 0.725 | 1.435 | 0.151 |
| **factor(Conditions)ASYNCH3PP × ERS** | 0.777 | 1.516 | 0.130 |

**Table S30.**

**Effect of hippocampal ERS and conditions on performance for recognition of changed scene (Change)**: Performance (Change) ~ Hippocampal ERS *Conditions +Trials + random(Participants).

| | Estimate | z-value | p-value |
|---|---|---|---|
| **(Intercept)** | 0.324 | 1.487 | 0.137 |
| **factor(Conditions)ASYNCH1PP** | 0.276 | 1.587 | 0.112 |
| **factor(Conditions)ASYNCH3PP** | 0.180 | 1.065 | 0.287 |
| **ERS** | 1.545 | 3.527 | <0.001*** |
| **Trials** | 0.052 | 8.290 | <0.001*** |
| **factor(Conditions)ASYNCH1PP × ERS** | -0.593 | -0.916 | 0.360 |
| **factor(Conditions)ASYNCH3PP × ERS** | -1.665 | -2.551 | 0.011 * |

**Table S31.**

**Effect of hippocampal ERS and conditions on performance for recognition of original scene (No change)**: Performance (NoChange) ~ Hippocampal ERS *Conditions +Trials + random(Participants).

| | Estimate | z-value | p-value |
|---|---|---|---|
| **(Intercept)** | -0.013 | -0.042 | 0.966 |
| **ERS** | 1.720 | 3.520 | <0.001*** |
| **Trials** | 0.075 | 6.260 | <0.001*** |

**Table S32.**

**Effect of hippocampal ERS on performance for recognition of original scene (No change) in SYNCH1PP**: Performance (No Change) ~ Hippocampal ERS SYNCH1PP +Trials + random(Participants).

| | Estimate | z-value | p-value |
|---|---|---|---|
| **(Intercept)** | 0.742 | 2.512 | 0.012 * |
| **ERS** | -0.223 | -0.429 | 0.668 |
| **Trials** | 0.043 | 4.036 | <0.001*** |

**Table S33.**

**Effect of hippocampal ERS on performance for recognition of original scene (No change) in ASYNCH3PP**: Performance (No Change) ~ Hippocampal ERS ASYNCH3PP +Trials + random(Participants).

**Movie S1.**

Supplementary Video 1

Experimental design: (Upper panel) 3D scene with the embedded avatar, as observed by a participant performing the virtual reality task. (Lower panel) Movements performed by a participant inside the MR scanner. Due to physical constraint (MR scanner magnetic field), this was filmed in a replicate of an MR scanner, similar to the one used for Experiment 1 and 3.
